# ppGpp influences protein protection, growth and photosynthesis in *Phaeodactylum tricornutum*

**DOI:** 10.1101/2020.03.05.978130

**Authors:** Luisana Avilan, Regine Lebrun, Carine Puppo, Sylvie Citerne, Stephane Cuiné, Yonghua Li-Beisson, Benoit Menand, Ben Field, Brigitte Gontero

## Abstract

- Chloroplasts retain elements of a bacterial stress response pathway that is mediated by the signalling nucleotides guanosine penta- and tetraphosphate, or (p)ppGpp. In the model flowering plant Arabidopsis, ppGpp acts as a potent regulator of plastid gene expression and influences photosynthesis, plant growth and development. However, little is known about ppGpp metabolism or its evolution in other photosynthetic eukaryotes.
- Here, we studied the function of ppGpp in the diatom *P. tricornutum* using transgenic lines containing an inducible system for ppGpp accumulation. We used these lines to investigate the effects of ppGpp on growth, photosynthesis, lipid metabolism and protein expression.
- We demonstrate that ppGpp accumulation reduces photosynthetic capacity and promotes a quiescent-like state with reduced proliferation and ageing. Strikingly, using non-targeted proteomics, we discovered that ppGpp accumulation also leads to the coordinated upregulation of a protein protection response in multiple cellular compartments.
- Our findings highlight the importance of ppGpp as a fundamental regulator of chloroplast function across different domains of life, and lead to new questions about the molecular mechanisms and roles of (p)ppGpp signalling in photosynthetic eukaryotes.

## Introduction

Diatoms are a group of unicellular eukaryotic photosynthetic organisms that form a major part of phytoplankton, and are responsible for up to one fifth of net global carbon fixation (Falkowski *et al.*, 2000). Beside their ecological importance, diatoms are also studied because they have a wide range of potential applications that include drug delivery in chemotherapy, biofuels, and as environmental indicators for monitoring water quality (Levitan *et al.*, 2014; Lavoie *et al.*, 2017; Uthappa *et al.*, 2018).

Diatoms appeared relatively recently in evolutionary history, around 200 million years ago (Medlin, 2016; Benoiston *et al.*, 2017). The diatom chloroplast was acquired through complex endosymbiotic events, where it is thought that a red algal ancestor was engulfed by a eukaryote that already possessed green algal genes from a previous endosymbiosis (Dorrell *et al.*, 2017). The acquisition of additional genes by lateral gene transfer from bacteria is also likely to have been an important driving force in the evolution of diatoms (Bowler *et al.*, 2008). These complex origins confer a unique cellular physiology to diatoms that allows them to adapt to multiple environments (Vardi *et al.*, 2008; Gruber & Kroth, 2017).

The diatom chloroplast differs in a number of aspects from the chloroplast of plants and green algae. At the level of membrane architecture, the diatom chloroplast is surrounded by four membranes, and the thylakoids are loosely stacked with three interconnected membranes (Flori *et al.*, 2017). Moreover, the light-harvesting complex (LHC) of diatoms is known as the Fucoxanthin Chlorophyll Protein complex (FCP), and is composed of tetramers containing chlorophyll c and the carotenoid fucoxanthin (Lepetit *et al.*, 2012; Roding *et al.*, 2018; Nagao *et al.*, 2019; Pi *et al.*, 2019). Diatoms have other specific features such as a functional urea cycle (Allen *et al.*, 2011), a eukaryotic Entner-Doudoroff glycolytic pathway (Fabris *et al.*, 2012) and they lack the plastid oxidative pentose phosphate pathway found in plants (Gruber & Kroth, 2017). The regulation of diatom metabolism is also distinct from that in plants (Jensen *et al.*, 2017).

Like plants, diatoms possess signalling pathways with prokaryotic origins such as the bacterial histidine-kinase-based two-component systems (Bowler *et al.*, 2008), as well as pathways of eukaryotic origins such as the Target Of Rapamycin (TOR) kinase (Prioretti *et al.*, 2017; Prioretti *et al.*, 2019) and the G protein-coupled receptor signalling pathway (Port *et al.* 2013).

Another signalling pathway that could play an important role in diatom stress acclimation is the pathway mediated by the nucleotides guanosine tetraphosphate and guanosine pentaphosphate or (p)ppGpp (Field, 2018; Avilan *et al.*, 2019). In bacteria, ppGpp and to a lesser extent pppGpp accumulate in response to a range of different stresses, and specifically target transcription and translation to slow growth and promote stress acclimation (Hauryliuk *et al.*, 2015; Steinchen & Bange, 2016). (p)ppGpp is also found in plants and green algae (Takahashi *et al.*, 2004), and chloroplast-localized RelA SpoT Homolog (RSH) enzymes for (p)ppGpp synthesis are widespread among the photosynthetic eukaryotes (Atkinson *et al.*, 2011; Ito *et al.*, 2017; Avilan *et al.*, 2019). Currently, the function of ppGpp is best characterised in the flowering plant Arabidopsis, where it inhibits chloroplast gene expression, reduces chloroplast size, and reduces photosynthetic capacity (Maekawa *et al.*, 2015; Sugliani *et al.*, 2016; Honoki *et al.*, 2018). While the mechanisms are still uncertain, ppGpp may act by downregulating the transcription of chloroplast encoded genes via the inhibition of chloroplastic RNA polymerases or GTP biosynthesis (Nomura *et al.*, 2014; Yamburenko *et al.*, 2015; Sugliani *et al.*, 2016). Levels of ppGpp increase in response to different abiotic stresses, as well as in response to treatment with stress-associated plant hormones (Takahashi *et al.*, 2004). Increased ppGpp levels can affect growth under nitrogen limiting conditions (Maekawa *et al.*, 2015; Honoki *et al.*, 2018) and plant immune signalling (Abdelkefi *et al.*, 2018).

Genes coding for RSH enzymes are present in all fully sequenced photosynthetic eukaryotes (Field, 2018; Avilan *et al.*, 2019). However, very little is known about the role of (p)ppGpp in algae, and in particular in algae of the red lineage. A recent study in the extremophile red alga *Cyanidioschyzon merolae* showed that overexpression of CmRSH4b, a functional (p)ppGpp synthetase, results in a reduction in chloroplast size and decreased chloroplast rRNA transcription, although (p)ppGpp levels were not measured (Imamura *et al.*, 2018). The situation in diatoms, with their mixed red-green heritage and specific lifestyles, is even less clear. A recent analysis showed that the nuclear genome of the marine diatom *Phaeodactylum tricornutum* encodes three functional RSH enzymes from red-lineage specific clades: PtRSH1, a bifunctional (p)ppGpp synthetase/hydrolase; and PtRSH4a and PtRSH4b, which act exclusively as (p)ppGpp synthetases (Avilan *et al.*, 2019). Here, we studied (p)ppGpp signalling in *P. tricornutum* by manipulating endogenous ppGpp levels using transgenic lines that express a heterologous (p)ppGpp synthetase under the control of an inducible promoter.

We found that ppGpp accumulation led to a reduction in photosynthetic capacity and an inhibition of ageing and growth. Strikingly, a proteomic analysis revealed that ppGpp accumulation also leads to the robust activation of a protein protection response involving chaperones and proteases. Our findings demonstrate that core ppGpp signalling is highly conserved across the photosynthetic eukaryotes, and that ppGpp has species specific roles that may be linked to adaptation to particular environments and life-styles.

## Materials and methods

### Cell culture

*P. tricornutum* Bohlin (strain name Pt1_8.6; RCC 2967) was obtained from Roscoff Culture Collection. Cells were grown in f/2 medium (Guillard & Ryther, 1962) without silica and adjusted to pH 8. The medium contained either 0.88 mM NaNO_3_ (f/2-NO_3_) or NH_4_Cl (f/2-NH_4_) as a nitrogen source. Cultures, in liquid medium in Erlenmeyer flask or on agar plates (2% bacto-agar), were maintained at 18 °C under continuous illumination (30 μmol photon m^−2^s^−1^) with LED lamps. Samples for cell counting (Malassez counting chamber) and protein extraction were withdrawn from liquid cultures.

### Plasmid construction and transformation of *P. tricornutum*

The vector pPha-NR carrying the nitrate reductase promoter and the *ble* gene for zeocin resistance (Chu *et al.*, 2016) (GenBank accession number JN180663, kindly provided by Pr. P. Kroth), was used as backbone for all plasmid constructions. The DNA fragments used for gene fusion and cloning in the final constructs were amplified by PCR using Q5 DNA polymerase (New England Biolabs) using standard conditions. The primers are specified in Table S1. The amplified fragments were assembled in the vector, previously digested with *Eco*RI and *Hind*III, by Sequence and Ligation Independent Cloning (SLIC) method (Jeong *et al.*, 2012).

The genes coding for a (p)ppGpp synthetase (SYN) and an inactive mutant form (D275G) of this synthetase (SYN^D>G^) were obtained by PCR, using plasmids described in Sugliani *et al.,* (2016) as templates. SYN corresponds to the (p)ppGpp synthetase Rel A from *E. coli* (residue 1 to 386). To target the enzymes to the chloroplast, the genes were fused by PCR to the sequence coding for the bipartite targeting sequence of the chloroplastic gamma ATP synthetase of *P. tricornutum* (GeneBank accession number U29898) whose ability to target proteins to the chloroplast is well characterised (Apt *et al.*, 2002). A Kozak sequence (AAG) was included in the primer before the ATG translation start site to facilitate translation. The final constructions were used to transform cells by particle bombardment.

Transformation of *P. tricornutum* was performed using a Bio-Rad Biolistic PDS-1000/He particle delivery system with rupture disc of 1350 psi as previously described (Falciatore *et al.*, 1999; Kroth, 2007). Prior to the bombardment, 1 × 10^7^ cells, in the exponential growth phase, were spread in the center (5 cm diameter) of the f/2-NH4 agar plate without antibiotic, dried under sterile hood and placed under illumination for 24 h. Ten μl of M17 tungsten microcarriers (1.1 μm diameter, Bio-Rad) equivalent to 600 μg particle were coated with 1 μg of the specific plasmid constructs in the presence of CaCl_2_ and spermidine (Kroth, 2007) and used to bombard the diatom cells. After 24 h illumination cells were resuspended and re-plated onto f/2-NH_4_ agar plates containing 80 μg mL^−1^ zeocin (Invitrogen). The plates were maintained under illumination for 3 weeks and the resistant colonies were screened by PCR for the presence of *SYN* genes as previously described (Falciatore *et al.*, 1999) (primers in Table S1). A cell suspension from each colony was re-plated on agar medium and the screening process was repeated. The final colonies were also screened for the presence of the proteins SYN by Western blotting. Interestingly, we obtained fewer SYN lines (n=9) than SYN^D>G^ (n=66), suggesting that the presence of SYN negatively affects the recovery of transgenics. Clones of *SYN* lines were prone to losing their phenotype after several multiplications. The transformants were therefore periodically re-plated from cells stored at 4°C and re-screened based on their growth phenotype on solid agar plates. Multiple independent transgenics lines were used for each construction in subsequent experiments.

### Nucleotide determination

GTP and ppGpp levels were determined using stable isotope labelled internal standards as described (Bartoli *et al.*, 2019). Briefly, 8 mg (dry weight equivalent) of *P. tricornutum* cells were harvested from cultures 2 days after induction and nucleotides directly extracted with 3 ml 2M formic acid containing 12.5 pmoles ^13^C-labelled ppGpp and 125 pmoles ^13^C labelled GTP. Extracts were incubated on ice for 30 minutes, 3 ml 50 mM ammonium acetate at pH 4.5 was then added, and then samples were loaded onto pre-equilibrated Oasis WAX SPE cartridges and nucleotides eluted with a mixture of methanol/water/NH_4_OH (20:70:10). The eluates were lyophilized and resuspended in water before analysis by HPLC-MS/MS with multiple reaction monitoring. The quantification of (p)ppGpp in Figure 1C was hindered by poor detection of the ^13^C-ppGpp internal standard during HPLC-MS/MS analysis. ppGpp concentrations were therefore calculated using the ^13^C-GTP internal standard adjusted for the response factor to estimate recovery. Figure S1 shows absolute quantification of ppGpp based on the ^13^C-ppGpp internal standard.

**Fig. 1.**
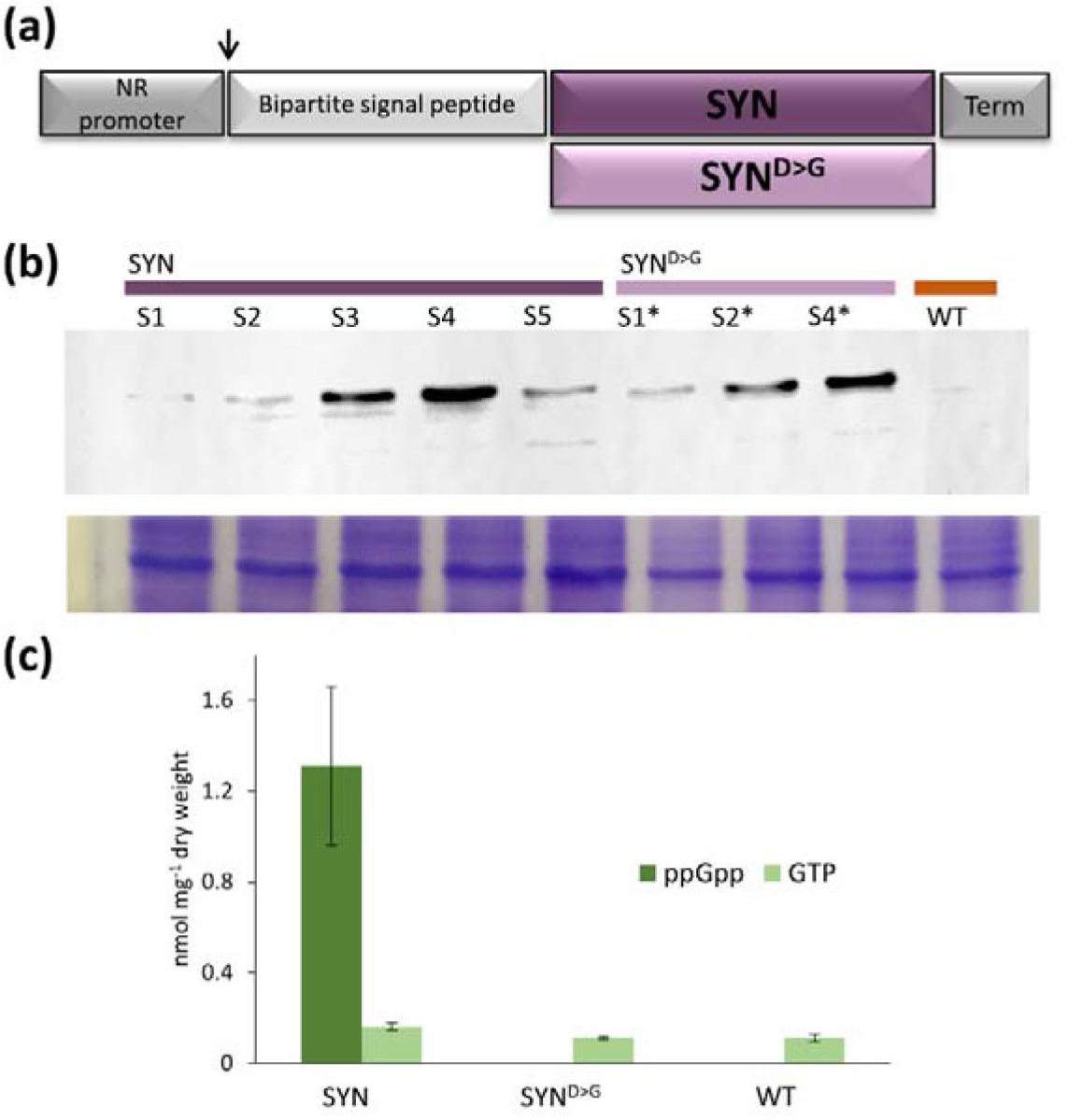
The creation of NO_3_ inducible lines for the accumulation of ppGpp in *P. tricornutum*. **(a)** Schematic representation of the construction used for the transformation of *P. tricornutum* cells. Nitrate reductase (NR) promoter and terminator (Term) were from pPha-NR vector. The bipartite targeting sequence was from the chloroplastic gamma ATP synthase. The genes encode a (p)ppGpp synthetase (SYN) and an inactive mutant form of SYN (SYN^D>G^). **(b)** Western blot analysis of extracts from different transgenic lines induced on f/2-NO_3_ using anti-*E. coli* RelA antibodies for SYN and SYN^D>G^. Each well was loaded with 7 ×10^4^ cells. Coomassie blue staining is shown as a loading control. **(c)** ppGpp and GTP levels in SYN (±SE, n=5 independent lines), SYN^D>G^ (±SE, n =4 independent lines) and WT lines (±SE, n=3 independent replicates).

### Growth parameters and pigment quantification

For the analysis of growth phenotypes cells were grown in 25 ml liquid f/2-NH_4_ in 250 ml Erlenmeyer flasks, without agitation, in triplicate for five different transformants. Optical density (OD) at 750 nm and number of cells were monitored every 24 hours. Cell number showed a linear relationship with OD 750 nm up to an OD of 1. Cultures were inoculated from a preculture to an initial OD 750 nm of 0.05. After 48 hours, cells were centrifuged at 5000 g (Allegra® X-15R Centrifuge, Beckman Coulter) for 10 minutes at 16 °C and the medium was substituted with f/2-NO_3_ to induce the expression of SYN or SYN^D>G^. For control cultures, the medium was substituted with fresh f/2-NH_4_. Pigments were extracted on ice from cell pellets (1.6 × 10^7^ cells) with 1 ml 96% (v/v) ethanol, maintained in the dark for 30 minutes and centrifuged at 12.000 rpm for 10 minutes at 4°C. The spectra of the supernatant from 350 to 750 nm were recorded using a Perkin Elmer spectrophotometer (PTP-6 Peltier System). Chlorophyll a and c concentrations were calculated according to (Ritchie, 2006) and fucoxanthin concentration according to (Wang *et al.*, 2018). For extended darkness, 24 hours after induction cells were transferred to darkness for 21 days.

### Light microscopy and cell measurement

Light microscope images of live cells were taken using a MoticBA410 microscope equipped with a Moticam 1080 camera. Images were used to determine length of the individual cells using the Motic image 3 plus software after calibration of the microscope. For fluorescence microscopy, cells were incubated with the fluorescent dye AC202 as described in (Harchouni *et al.*, 2018). Fluorescence was visualized in an epifluorescence microscope Eclipse 80i (Nikon) using the DAPI filter cube (excitation filter: 360BP40; emission filter 460BP50). Images were acquired using MS Elements imaging software (Nikon). Images were merged using ImageJ software.

### Measurement of photosynthetic activity

Chlorophyll fluorescence parameters were measured using a Fluorcam FC 800-O imaging fluorometer (Photon System Instruments, Drasov, Czech Republic). Cells plated on solid medium were recovered in f2/NH_4_ medium and 10 μl of this cell suspension (between 5 and 9 × 10^7^ cells mL^−1^) was dotted onto f2/NO_3_ or f2/NH_4_ medium and grown under standard growth conditions. Before measurement, the plates were kept in dark for 20 minutes and the chlorophyll fluorescence imaged to obtain minimum fluorescence yield (F_o_) and maximum fluorescence yield (F_m_). PSII maximum efficiency was calculated as F_v_/F_m_ = (F_m_−F_o_)/F_m_. To calculate relative electron transfer rate (ETR), samples were exposed to different photon flux densities (PPFD) in a stepwise fashion. Relative ETR was then calculated as the product of the photochemical yield of PSII (ΦP = ΔF/F_m_′ = (F_m_′−F_o_)/F_m_′) and PPFD.

### Cell extract, SDS-PAGE and immunoblotting

*P. tricornutum* cells (2 × 10^7^) were harvested from liquid culture by centrifugation (5000 g for 15 minutes at 18°C) and resuspended in 100 μl of rupture buffer (10 mM Tris, pH 8 containing 2% n-dodecyl β-D-maltoside) and then broken by sonication using 6 pulses using an ultrasonicator (Sonics & Materials Inc, Vibracell, Bioblock, Danbury, Connecticut, USA). After incubation at 4°C for 30 minutes, the cell extract was centrifuged at 12000 g for 30 minutes at 4°C. The supernatant was submitted to SDS-PAGE and gels were stained with Coomassie blue R-250. For Western blot analysis, proteins were transferred onto nitrocellulose membrane as described (Sambrook *et al.*, 1989). Loading control gels were run in parallel. The membrane was blocked with TBS (Tris 10 mM, pH 8, 150 mM NaCl) containing 5% nonfat milk and incubated with the specified primary antibodies in the same buffer containing 1% nonfat milk. The antibodies used were: anti-*E.coli* RelA (dilution 1/2000, kindly provided by M. Cashel), anti-hemagglutinin (HA) (monoclonal, dilution 1/10000, Sigma-Aldrich, catalog number:H9658, clone HA-7), anti-PsbA (polyclonal, dilution 1/10000, Agrisera, catalog number AS05 084) and anti-*P. tricornutum* LHCf1-LHCf11 (Juhas and Buchel, 2012) (polyclonal, dilution 1:5000, kindly provided by C. Büchel). After washing with TBS, the membranes were incubated either with horseradish peroxidase coupled anti-rabbit IgG (GE Healthcare, catalog number NA934V) for the polyclonal antibodies or anti-mouse IgG (Sigma-Aldrich) for the monoclonal antibodies. Immunodetection was performed using the enhanced chemiluminescence method (ECL substrates, GE Healthcare).

### Lipid and chrysolaminarin determination

The fluorescent probe Nile red was used to detect neutral lipids by flow cytometry as specified in (Prioretti *et al.*, 2017). The cells at different growth phases were fixed with 2% (v/v) glutaraldehyde (Prioretti *et al.*, 2017) and 1 ml of cell suspension was incubated with 2 μl of Nile red solution (0.25 mg ml^−1^) for 5 minutes before the analysis with a bench flow cytometer (BD Accuri C6, BD Biosciences). To determine Nile red fluorescence, the trigger signal was set to FL2 fluorescence and combined with the side scatter (SSC) signal. Ten thousand events were analysed in each case.

For lipid analysis, cells of *P. tricornutum* (50 × 10^6^ cells) were harvested by centrifugation (5000 *g*, for 10 minutes) and the pellet was resuspended with 1 mL of hot isopropanol (preheated at 85°C) and incubated at this temperature for 10 minutes to quench endogenous lipases. Detailed lipid extraction protocol can be found in (Legeret *et al.*, 2016). Extracted total lipids were then dried under a stream of N_2_, and then dissolved in a solvent mixture of chloroform: methanol (2:1 vol/vol) for quantification by Thin Layer Chromatography (TLC). Two types of TLC were run: one for triacylglycerol (TAG) quantification, and a second for polar membrane lipid analysis. Detailed TLC procedures, standard used and quantifications have been described, together with total fatty acid (FA) content and composition analysis in (Siaut *et al.*, 2011).

Extraction and determination of the β-1,3-glucan chrysolaminarin was performed following the method of (Granum & Myklestad, 2002) from pellets containing 2 × 10^7^ cells. A calibration curve was obtained using glucose as a standard.

### Mass spectrometry analysis

Proteins extracts (50 μg protein) from 3 SYN lines, 2 SYN^D>G^ lines and 1 WT, made on equal number of cells, were loaded on a gel (SDS-PAGE) and the band from the stacking gel corresponding to the total proteins was excised and submitted to in gel trypsin digestion for proteomic analysis as previously described in (Santin *et al.*, 2018) with minor modifications. In parallel, total proteins were also separated to analyse the protein profile in the gel (Fig. S1). Tryptic peptide samples were quantified by a colorimetric peptide assay (Pierce, Thermo Fisher) and aliquots of 200 ng were injected by LC-MS/MS. Two LC-MS/MS injections were performed per condition (technical replicates). Peptides were separated on an analytical C18 column by a two step -linear gradient from 4% to 20% of mobile phase B (0.1% (vol/vol) formic acid (FA)/ 80% (vol/vol) acetonitrile) in mobile phase A (0.1% (vol/vol) FA) for 90 minutes, then from 20% to 45% of B in A for 30 minutes. For peptide ionization in the nanosource spray, voltage was set at 1.65◻kV and the capillary temperature at 275◻°C. Top 10 Data Dependent workflow was used in a 400-1600 m/z range and a dynamic exclusion of 60 s.

Spectral data were processed for protein identification and quantification using the MaxQuant computational proteomics platform (version 1.6.5.0) integrating the search engine Andromeda and the MaxLFQ algorithm (Cox *et al.*, 2014). Spectra were searched against a UniProt *P. tricornutum* database (date 2018.01; 10715 entries) supplemented with a set of 245 frequently observed contaminants. The search parameters were set at: (*i*) trypsin cleavage authorized before proline with two missed cleavages allowed; (*ii*) monoisotopic precursor tolerance of 20 ppm in the first search used for recalibration, followed by 4.5 ppm for the main search and 0.5 Da for fragment ions from MS/MS; (*iii*) cysteine carbamidomethylation (+57.02146) as a fixed modification and methionine oxidation (+15.99491) and N-terminal acetylation (+42.0106) as variable modifications; (*iv*) a maximum of five modifications per peptide allowed and (*v*) minimum peptide length was 7 amino acids and a maximum mass of 4,600 Da. The false discovery rate (FDR) at the peptide and protein levels were set to 1% and determined by searching a reverse database. Statistical analysis was performed with Perseus (version 1.5.6.0) (Tyanova *et al.*, 2016). LFQ normalized intensities were uploaded from the proteinGroups.txt and converted to base 2 logarithms to obtain a normal distribution. Quantifiable proteins were defined as those detected in at least 70% of samples in at least one condition. Missing values were replaced using data imputation by randomly selecting from a normal distribution centred on the lower edge of the intensity values. To determine whether a given detected protein showed differential accumulation, a two-sample t-test was applied using a permutation based FDR-set at a conservative threshold of 0.0001 (250 permutations), and the p value was adjusted using a scaling factor s0 set to 1. The results are illustrated in a volcano plot (Fig. S2), and the list of identified proteins and differentially accumulating proteins in SYN versus the control lines is available in Table S2.

## Results

### An inducible (p)ppGpp synthetase efficiently increases ppGpp levels in *P. tricornutum*

As a first step towards understanding the function of (p)ppGpp in diatoms, we developed a strategy similar to that reported by (Sugliani *et al.*, 2016) to modulate (p)ppGpp levels in the chloroplast. For this purpose, we transformed *P. tricornutum* with a synthetic gene encoding a chloroplast-targeted fragment of a bacterial (p)ppGpp synthetase (SYN) under the control of an NO_3_ inducible promoter (Fig. 1a). Control transgenic lines were also generated that express catalytically inactive forms of the same enzyme, SYN^D>G^. Positive transformants were identified by PCR and analysed to confirm protein expression following induction by transfer from NH_4_-containing medium to NO_3_-containing medium (Fig. 1b). Importantly, induction of SYN lines also caused an increase in ppGpp levels to 1.30 ± 0.34 nmol mg^−1^ dry cell weight (n=5 independent biological replicates) (Fig. 1c). pppGpp was not detected, suggesting that SYN preferentially acts as a ppGpp synthetase, and/or that pppGpp can be converted to ppGpp by endogenous enzymes as in bacteria. No ppGpp was detected in SYN^D>G^ lines or the wild type under inducing conditions. The absence of basal levels of ppGpp is likely to be a consequence of the nutrient shift used for induction because we were able to detect low quantities of ppGpp in wild type cells grown in standard media in the light (Fig. S1). Interestingly, we also observed that incubation of cells in the dark for prolonged periods led to the accumulation of ppGpp. We simultaneously quantified GTP levels and found that concentrations were similar to the wild-type control regardless of ppGpp levels (Fig. 1c, Fig. S1).

### ppGpp accumulation strongly inhibits cell division and has a major effect on photosynthesis

We first examined the effects of ppGpp accumulation on growth and proliferation. Five independent SYN lines showed a severe reduction in proliferation upon induction in liquid culture (Fig. 2a and Fig. S2). In contrast, induction did not affect the proliferation of control SYN^D>G^ lines or the WT. The reduced proliferation of SYN lines was also clearly visible on agar plates following induction (Fig. 2b). Observation by light microscopy showed that SYN cells were significantly longer than those of the controls (Fig. 2c).

**Fig. 2.**
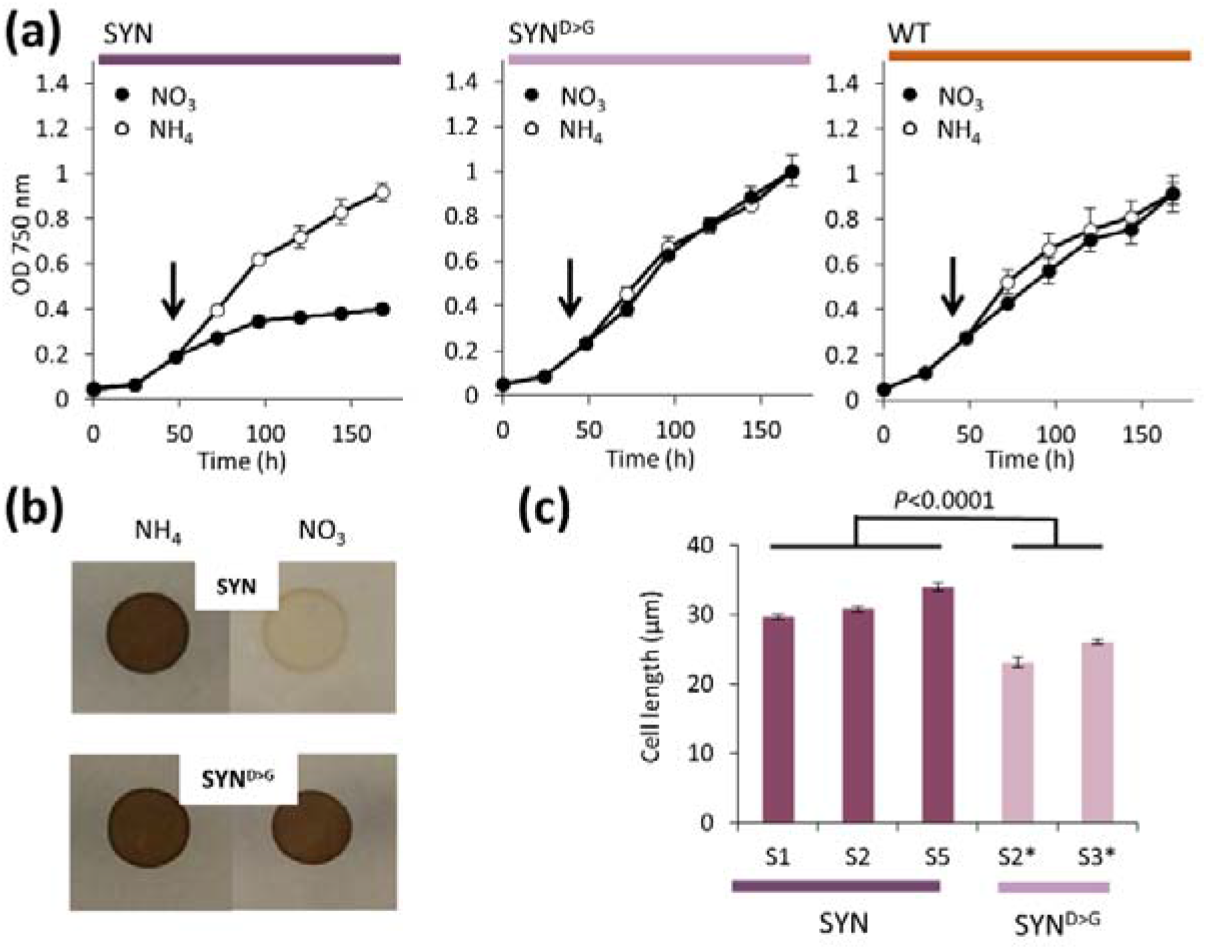
ppGpp levels affect proliferation and cell size. **(a)** Growth curves of SYN, SYN^D>G^ and WT lines. Cells grown in f/2-NH_4_ were transferred (arrow) to either f/2-NO_3_ for induction or f/2-NH_4_ as controls. **(b)** Cells (SYN, SYN^D>G^) initially grown in liquid f/2-NH_4_ were grown as cell colonies on agar plates either in the presence of NO_3_ or in the presence of NH_4_ as controls. **(c)** Cell lengths of SYN and SYN^D>G^ cells after induction (±SE, n between 60 and 194 cells). Statistical significance was calculated using ANOVA with post-hoc Dunnett test.

We then analysed pigment content at different times after induction (Fig. 3, Fig. S3). In the WT, the level of chlorophyll a (Chl*a*) and fucoxanthin decreases as the cells enter stationary phase, five days post induction. To our surprise, we found that this drop in pigment content does not occur in induced SYN lines: five days after induction levels of Chl*a* and fucoxanthin are significantly higher in SYN than in SYN^D>G^ or WT (Fig. 3). In contrast, the drop in Chl*c* levels was similar in all lines, leading to a higher Chl*a*:Chl*c* ratio in the SYN lines (Fig. 3). These results suggest that ppGpp accumulation inhibits cell division while stabilizing Chl*a* and fucoxanthin levels in the chloroplast.

**Fig. 3.**
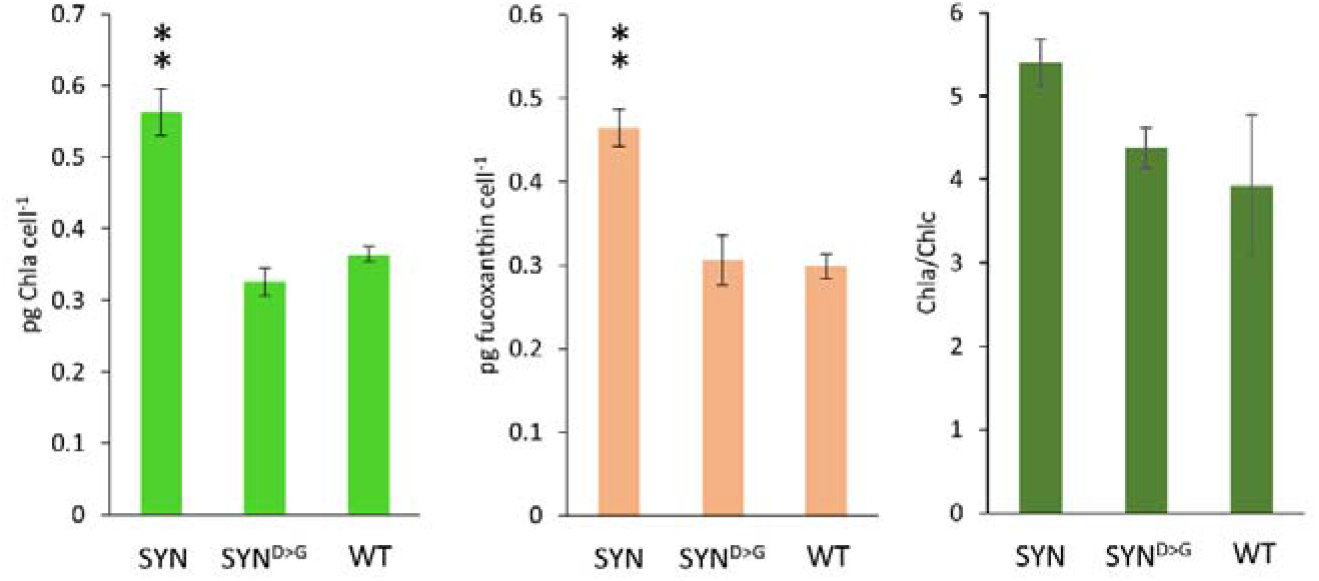
Pigment levels are maintained in induced SYN lines. Cells grown in f/2-NH4 were transferred to f/2-NO3 for induction. 5 days post induction, corresponding to the stationary phase of the growth curve, pigments were extracted with ethanol and the levels of chlorophyll a, chlorophyll c and fucoxanthin were determined. Data are means ± SE of five biological replicates. Statistical significance was calculated using ANOVA with post-hoc Dunnett tests versus WT control, **P < 0.01.

In Arabidopsis, ppGpp accumulation has major effects on photosynthetic activity (Maekawa *et al.*, 2015; Sugliani *et al.*, 2016). We therefore examined the maximal efficiency of photosystem II (F_v_/F_m_) (Fig. 4a-c). F_v_/F_m_ decreased rapidly in induced SYN transformants following induction, reaching a minimum of 0.12 ± 0.004 (SE) two days post induction (Fig. 4a,b and Fig. S4). The F_v_/F_m_ of the SYN^D>G^ lines was 0.6 at the same timepoint. We performed immunoblots to determine whether the decrease in F_v_/F_m_ in induced SYN lines was due to changes in PSII architecture. Indeed, we found that the levels of the chloroplast-encoded protein D1, that constitutes the core of the PSII reaction centre, decreased in SYN lines (Fig. 4d). In contrast, the levels of the light-harvesting protein, LHCf, that forms a major part of the FCP antenna, remained relatively constant (Fig. 4d). These results indicate that the composition of PSII undergoes major changes in response to ppGpp accumulation. In agreement with the reduced PSII efficiency, the relative electron transfer rate (rETR) of the entire photosynthetic chain was also lower in SYN lines than in the WT or SYN ^D>G^ controls (Fig. 4e,f).

**Fig. 4.**
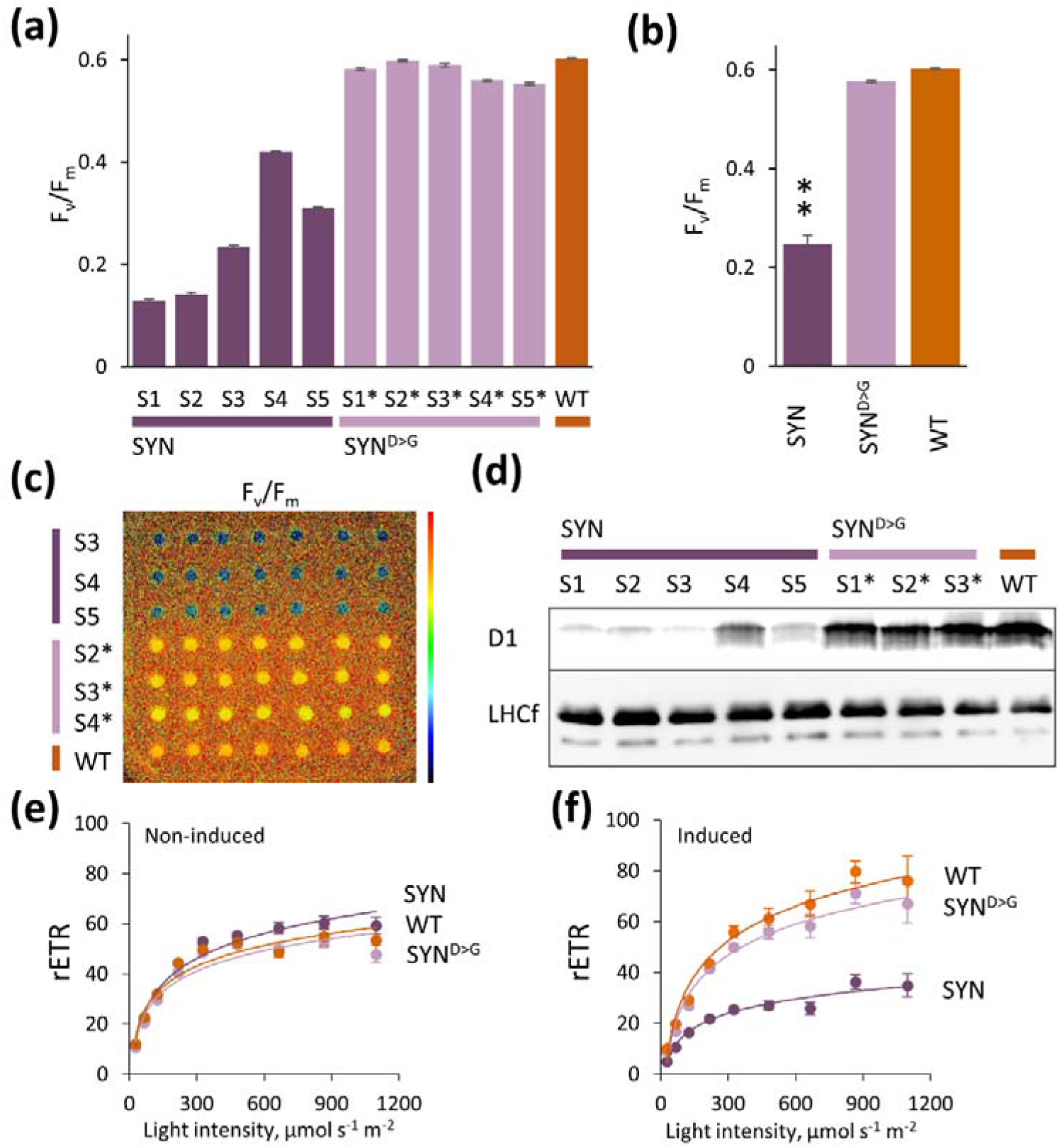
ppGpp accumulation downregulates photosynthetic activity. **(a)** The maximal efficiency of photosystem II (F_v_/F_m_) was measured on dark-adapted cells (n=7 colony spots) of different SYN, SYN^G>D^ and WT lines 2 days post induction. **(b)** Average F_v_/F_m_ of grouped lines for SYN, SYN^D>G^ and WT shown in A. **(c)** F_v_/F_m_ false colour image of cell colonies of SYN, SYN^D>G^ and WT one day post induction. **(d)** Immunoblots of protein extracts from equal number of cells (7 × 10^4^) two days post induction using primary antibodies against D1 and LHCf1-11. Relative electron transfer rate (rETR) at different light intensities in **(e)** uninduced and **(f)** induced SYN, SYN^D>G^ and WT lines two days post induction. Data are means ± SE of grouped lines for SYN and SYN^D>G^ (n=7 colony spots). Statistical significance was calculated using ANOVA with post-hoc Dunnett tests versus WT control, **P < 0.01.

### ppGpp affects the accumulation and distribution of lipids and other reserve compounds

Using light microscopy, we observed a prominent spot near, but clearly separated from, the chloroplast in all SYN lines at two days post induction (Fig. 5a). This spot was absent in non-induced SYN (Fig. 5b), and was not detected in SYN^D>G^ lines regardless of the condition (Fig. 5c, Fig. 5d). The prominent spot that was clearly visible under light microscopy was strongly stained by the neutral lipid specific fluorophore AC202, indicating that it is a lipid droplet (LD) (Fig. 5e). Moreover, AC202 staining also revealed the presence of small chloroplast associated LDs in the control SYN^D>G^ line (Fig. 5f) and the wild type. These droplets differ in size and location from the prominent cytoplasmic LD observed in induced SYN lines, and are not visible by light microscopy (Fig. 5e).-The prominent LD in SYN cells disappeared five days after induction (Fig. 5g). This corresponds to the stationary phase of the growth curve, when numerous LDs appear in SYN^D>G^ and WT (Fig. 5h). LDs are indeed well known to accumulate in diatoms with the ageing of the culture (Hu *et al.* 2008).

**Fig. 5.**
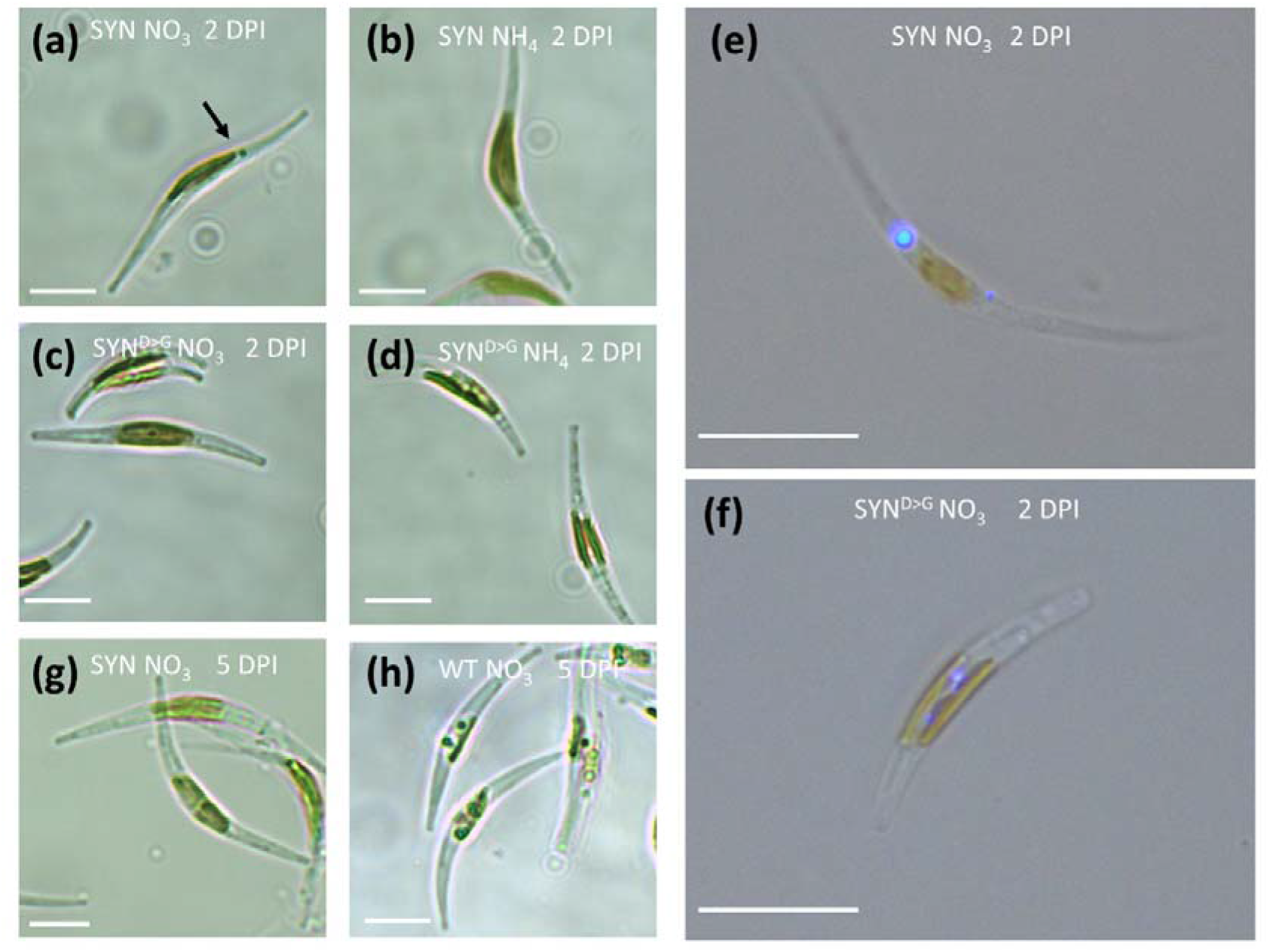
ppGpp accumulation affects lipid droplet formation. **(a)** Induced and **(b)** non-induced SYN cell after two days post induction (48 hr). **(c)** Induced and **(d)** non-induced SYN^D>G^ cells two days post induction. Fluorescence microscopy images of **(e)** SYN and **(f)** SYN^D>G^ cells two days post induction labelled with the fluorophore AC202. **(g)** SYN and **(h)** SYN^D>G^ cells five days after induction (120 hr). Images are representative of multiple images and at least two independent experimental replicates. The scale bar corresponds to 10 μm length.

To further characterize the prominent LD in SYN cells, we analysed neutral lipid content at different stages of the growth using flow cytometry and Nile red staining and then confirmed by quantification using TLC (Fig. 6a). A strong correlation between Nile red fluorescence and TAG content is established in diatoms (Greenspan *et al.*, 1985; Prioretti *et al.*, 2017). Nile red fluorescence, determined using flow cytometry, increased with the age of the culture in WT and SYN^D>G^ cells, but not in SYN cells, in agreement with our microscopic observations (Fig. 5g, h). However, flow cytometry was not sensitive enough to detect differences in lipid content at two days post induction. Therefore, we directly quantified TAG by extraction of total lipids followed by TLC quantification (Fig. 6b, c). Two days post induction, the amount of TAG per cell was almost two-fold higher in the SYN lines than in the controls (*P* =0.213) (Fig. 6b). Although not significant, this finding tends to support our microscopic observation of higher neutral lipid levels in SYN lines two days after induction (Fig. 5a). However, five days post induction, SYN lines contained much lower (6-fold) levels of neutral lipids than the control lines, in agreement with the flow cytometry results (Fig. 6c) and microscopy images (Fig. 5g, h). Chrysolaminarin (β-1,3-glucan), the main carbohydrate storage compound in diatoms which is located in the vacuole (Suzuki & Suzuki, 2013), was also less abundant in induced SYN lines than in the controls five days post induction (Fig. S6).

**Fig. 6.**
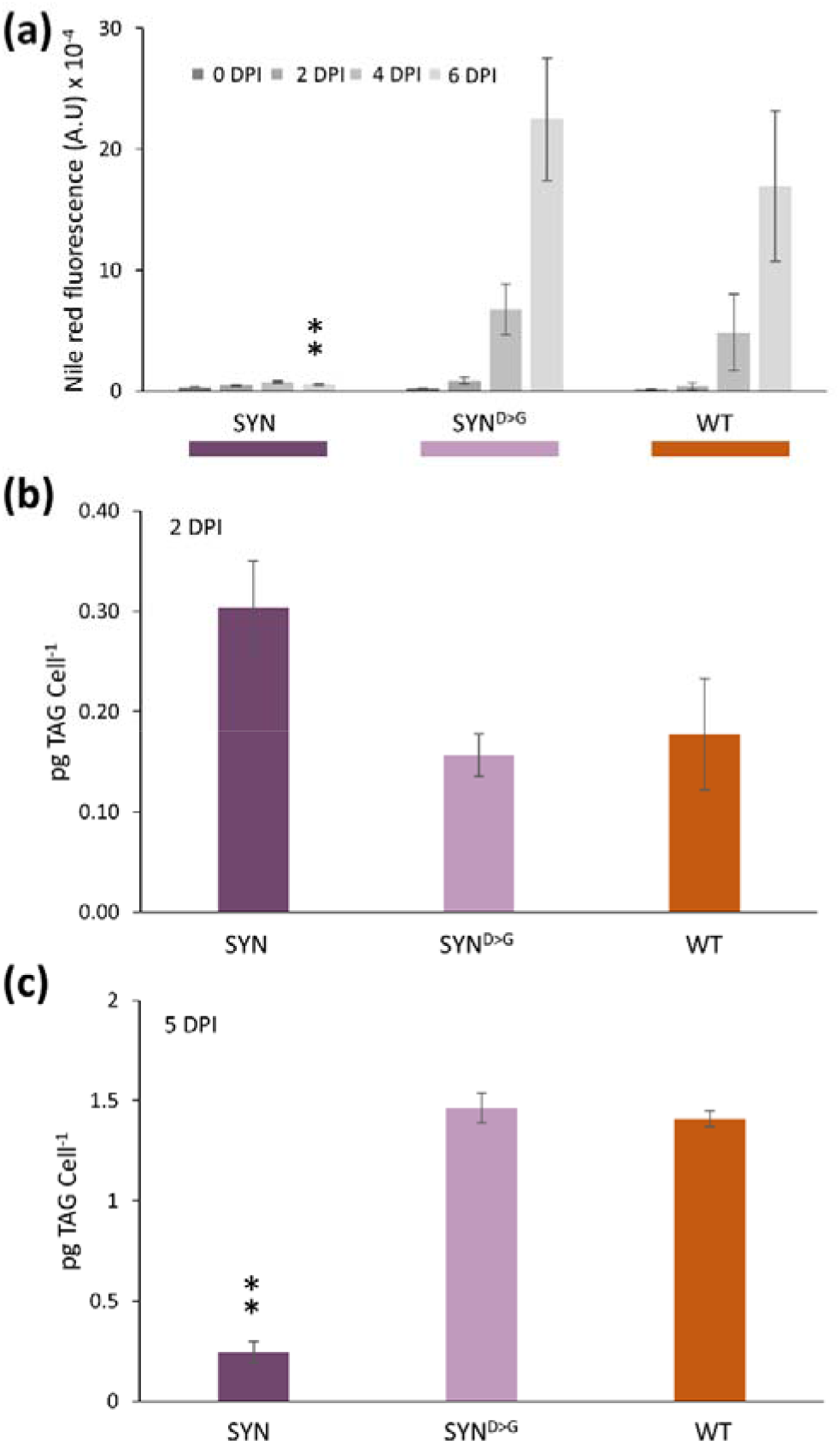
ppGpp accumulation inhibits neutral lipid accumulation in the stationary phase. **(a)** Neutral lipid estimation by Nile red fluorescence in SYN, SYND>G and WT lines at different days of the growth curve (inset). Data are means ± SE of three biological replicates. TAG determination by TLC **(b)** two days post induction and **(c)** five days post induction. Data are means ± SE of five biological replicates. Statistical significance was calculated using ANOVA with post-hoc Dunnett tests versus WT control, **P < 0.01.

To investigate the impact of altered ppGpp levels on membrane lipids, we also analysed polar lipid content. In the membranes of the diatom chloroplast, as in the chloroplasts of other photosynthetic organisms, polar lipids are essentially composed of monogalactosyldiacylglycerol (MGDG), digalactosyldiacylglycerol (DGDG), sulfoquinovosyldiacylglycerol (SQDG) and phosphatidylglycerol (PG) (Abida *et al.* 2015). Polar lipids in SYN cells two days after induction did not significantly differ from those of WT or SYN^D>G^ controls (Fig. S5). However, five days after induction, with the exception of DGDG and PG, the level of all the other polar lipids (MGDG; SQDG; phosphatidylcholine, PC; phosphatidylethanolamine, PE; phosphatidylserine, PS) remained higher in SYN cells than in the SYN^D>G^ and WT controls (Fig. 7a). Furthermore, FA content and composition in SYN cells was different compared to the controls five days after induction with substantially lower levels of the most abundant FAs (16:0 and 16:1), and decreases in 18:1, 18:2. These changes are consistent with the lower TAG levels in SYN lines, because 16:0, 16:1, 18:1 and 18:2 are the main FA species found in TAG under nutrient limiting conditions in *P. tricornutum* (Abida *et al.*, 2015).

**Fig. 7.**
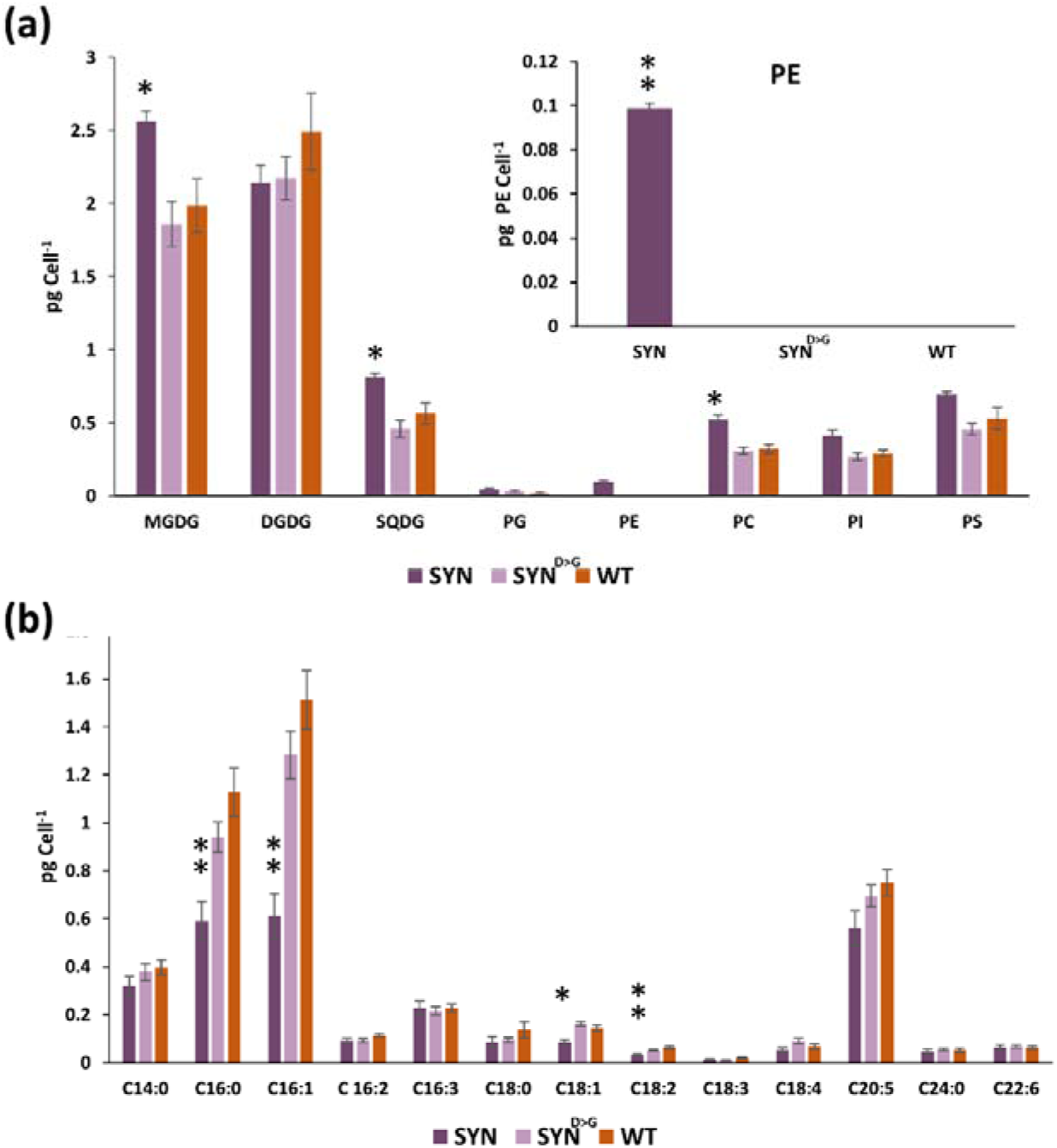
Effects of ppGpp on polar lipid and fatty acid composition. **(a)** Polar lipid and **(b)** fatty acid content were determined five days post induction. MGDG, monogalactosyldiacylglycerol; DGDG, digalactosyldiacylglycerol; SQDG, sulfoquinovosyldiacylglycerol; PG, phosphatidylglycerol; PE, phosphatidylethanolamine; PC, phosphatidylcholine; PI, phosphatidylinositol and PS, phosphatidylserine. Data are means ± SE of five biological replicates. Statistical significance was calculated using ANOVA with post-hoc Dunnett tests versus WT control, *P<0.05, **P < 0.01.

### Proteome analysis reveals that ppGpp accumulation leads to the coordinated upregulation of the protein protection response

The analysis of the protein profiles of induced SYN, SYN^D>G^ and WT by SDS-PAGE revealed major changes in total proteins in SYN lines, including the appearance of additional protein bands (Fig. S7). In order to understand the extent of these changes, we performed an untargeted proteomic analysis to compare protein accumulation between induced SYN lines and induced SYN^D>G^ lines two days after induction. Of the 1046 proteins that were identified, 145 proteins showed a significant difference in accumulation in SYN compared to the control lines (Table S1). The expression of known housekeeping proteins (Siaut *et al.*, 2007) was not affected by ppGpp accumulation, with the exception of actin 12 (Table S1). The major groups of proteins showing differential accumulation in SYN are listed in Table I. Unexpectedly, many chaperones and co-chaperones showed higher levels of expression in SYN, and the chaperone HSP20 (accession number B7G195) showed the greatest increase. The differentially expressed chaperones in SYN are encoded by nuclear genes and have different predicted cellular localizations. Four chaperones including HSP40 and one isoform of HSP90 contain the ASAFAP hexapeptide motif that predicts likely chloroplast localization (Gruber *et al.*, 2015). Other chaperones have signal peptides for the secretory pathway without an ASAFAP motif, suggesting localization in the endoplasmic reticulum. Some chaperones such as the Lon protease, that functions as both a chaperone and protease, are predicted to be mitochondrial.

We found that many transporters and enzymes involved in amino acid metabolism were among those proteins showing the greatest differential accumulation in the SYN lines (Table 1). Notably, the synthesis of glutamate appeared to be upregulated due to greater accumulation of the chloroplastic glutamate synthase and glutamate dehydrogenase, and lower levels of glutamine synthase. Glutamate is involved in nitrogen assimilation, and is also a precursor for chlorophyll biosynthesis. Interestingly, the chloroplast protoporphyrin IX magnesium chelatase, a key enzyme of the chlorophyll biosynthetic pathway, was also more abundant. Proteins of lipid metabolism were also affected. The levels of two enzymes involved in fatty acid degradation increased: the acyl-CoA dehydrogenase and a homologue of enoyl-CoA hydratase, a key enzyme of fatty acid beta-oxidation. Phage shock protein A, a protein that is involved in managing extra-cytoplasmic stress responses in bacteria (Joly *et al.*, 2010), and glutathione S-transferase, an enzyme important in the redox homeostasis of the cell were also more abundant.

**Table I.**
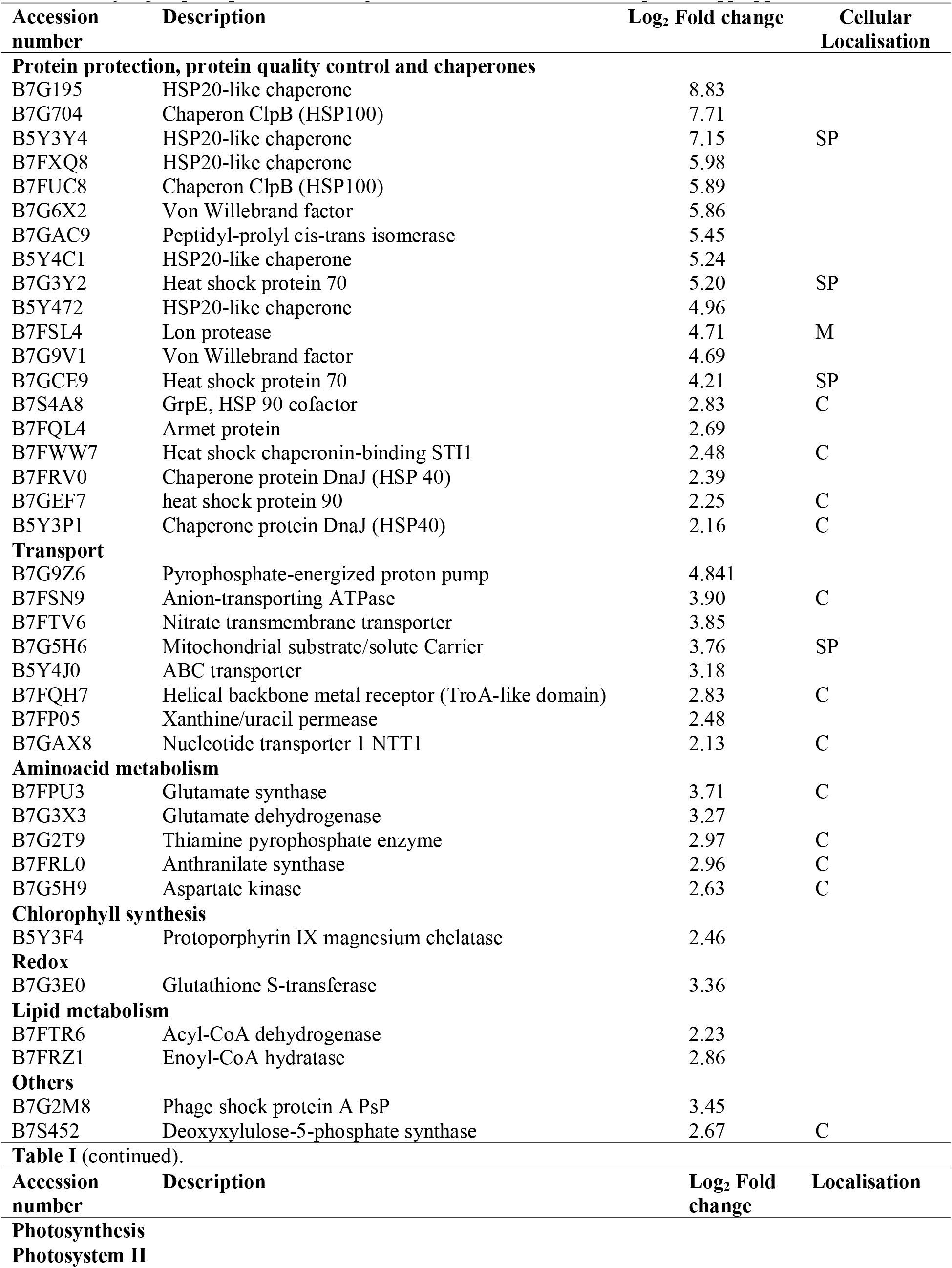
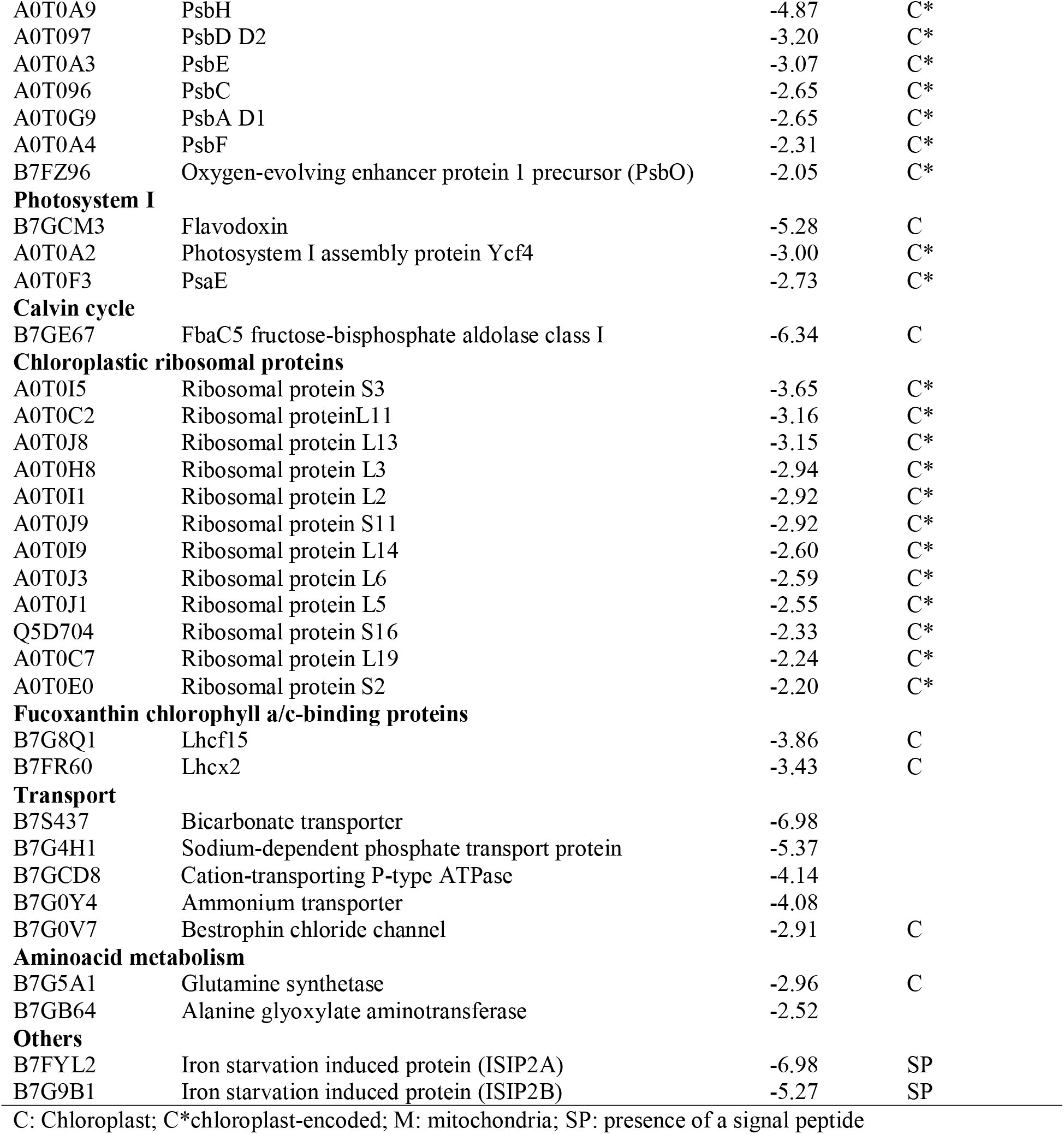
Major groups of proteins showing differential accumulation in response to ppGpp

Many chloroplast-encoded proteins with links to photosynthesis and chloroplast translation were less abundant in SYN. Seven proteins from the PSII complex accumulated to lower levels, including the PsbA / D1 protein, in agreement with our immunoblotting experiments (Fig. 4d). Some proteins from PSI and the Calvin cycle were also less abundant, as well as a bicarbonate transporter and two carbonic anhydrase proteins that were just outside the selection cut-off (Table S1), together indicating that photosynthetic capacity is diminished at several different levels. The photosynthetic machinery requires large amounts of iron, and consistent with the downregulation of photosynthesis we also observed the downregulation of two iron-starvation induced proteins. Interestingly, two light harvesting antenna proteins were also less abundant, Lhcf15 and Lhcx2. Therefore, although our immunoblotting experiment showed that Lhcf1-11 remained unchanged (Fig. 4d), SYN induction also appears to cause changes in the accumulation of specific antenna isoforms. Also striking was the reduced levels of twelve chloroplast-encoded ribosomal proteins in SYN cells, suggesting that ppGpp accumulation has a major effect on chloroplast translation capacity. The ribulose-1,5-bisphosphate carboxylase/oxygenase (Rubisco) remained unaltered in our proteomic analysis, as was also directly visible on Coomassie stained protein gels (Fig. S7).

## Discussion

In the present work, we studied the function of (p)ppGpp using transgenic *P. tricornutum* SYN lines containing an inducible system for the expression of a bacterial (p)ppGpp synthetase (Fig. 1a). We demonstrate that increased ppGpp levels in SYN lines alter the architecture of the photosynthetic machinery and reduce photosynthetic efficiency (Fig. 4). In addition, ppGpp accumulation dramatically decreases the growth rate (Fig. 2), stabilizes the level of chlorophyll and fucoxanthin (Fig. 4), and affects accumulation of the reserve molecules TAG and chrysolaminarin (Fig. 5–7, Fig. S6). Finally, ppGpp accumulation causes a striking increase in the levels of chaperones and other stress-related proteins in multiple cellular compartments (Table 1, Fig. 8).

**Fig. 8.**
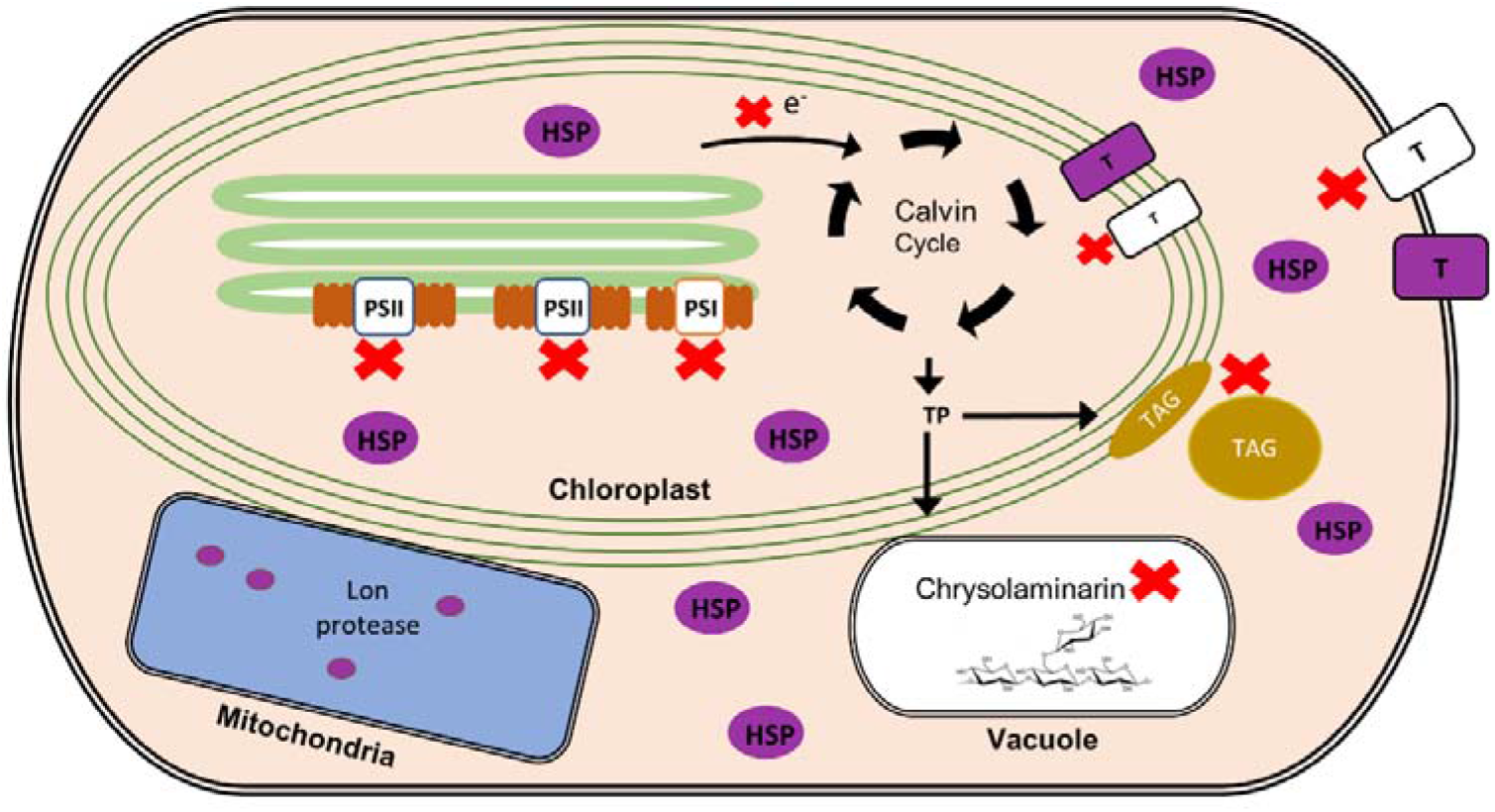
Summary of the effects of ppGpp accumulation on *P. tricornutum*. ppGpp accumulation causes a decrease in photosynthetic capacity, the chloroplast translation machinery, certain transporters, and the storage molecules TAG and chrysolaminarin (indicated by red X). At the same time, ppGpp accumulation causes a large increase in the abundance of proteins involved in the protein protection response (shown in purple). TP, triose phosphate; PSII and PSI, photosystems II and I; HSP, heat shock proteins; T, transporter.

The independent SYN lines produced different quantities of SYN and accumulated different quantities of ppGpp (Fig. 1). The differences we observed in SYN protein accumulation are likely due to differences in the copy number or the insertion site(s) of the exogenous DNA as previously shown (Falciatore *et al.*, 1999). The average levels of ppGpp reached in induced SYN lines (1.3 nmol mg^−1^ dry cell weight) are similar to those reached in *E. coli* (1.2 to 3 nmol mg^−1^ dry cell weight) during the stringent response (Riesenberg, 1985; Rodionov & Ishiguro, 1995). If we assume that total GTP content is similar between species, then these levels of ppGpp (representing 600% of total GTP) are considerably higher than those observed in Arabidopsis overexpressing a similar (p)ppGpp synthetase (9% of total GTP) (Sugliani *et al.*, 2016). This may be due to differences between unicellular and multicellular organisms, or the capacity of the cell to control ppGpp levels. Interestingly, the levels of GTP were very similar in all lines, indicating that ppGpp accumulation does not affect the total GTP cellular pool. This result resembles the situation in Arabidopsis ppGpp over-accumulating lines (Bartoli *et al.*, 2019) and contrasts with Gram-positive bacteria, where (p)ppGpp regulates transcription by decreasing the GTP pool in the cell (Krasny & Gourse, 2004).

We also detected ppGpp in WT cells and discovered conditions that cause ppGpp levels to increase (Fig. S1). Low ppGpp levels could be detected in WT cells under normal growth conditions. These ppGpp levels correspond to 0.25% of total GTP, which is similar to Arabidopsis where ppGpp accumulates to 0.36% of total GTP under normal growth conditions (Bartoli *et al.*, 2019). While nitrogen deprivation did not affect ppGpp levels, extended dark treatment caused a 4-fold increase in WT and SYN control lines (Fig. S1). Dark treatment also causes an increase in ppGpp levels in Arabidopsis in a CRSH dependent manner (Ihara *et al.*, 2015; Ono *et al.*, 2019). However, *P. tricornutum* lacks a direct orthologue of CRSH (Avilan *et al.*, 2019), suggesting differences in the underlying mechanism controlling (p)ppGpp synthesis in the dark.

In plants, ppGpp accumulation inhibits photosynthesis, in particular causing a large decrease in the abundance of Rubisco, and in the ratio of PSII reaction centre (RC) to PSII LHC (Maekawa *et al.*, 2015; Sugliani *et al.*, 2016). Similar results were obtained in this study (Fig. 4). We found that photosynthesis was strongly inhibited in induced SYN lines, and that this was accompanied by a large drop in the PSII RC to FCP antenna ratio and no obvious change in Rubisco levels (Fig. 4). Notably, these changes occur despite the significant differences in the structure and organisation of the diatom FCP antenna system compared to the equivalent light harvesting system in plants (Nagao *et al.*, 2019; Pi *et al.*, 2019). The overall rate of photosynthesis was also reduced by ppGpp accumulation (Fig. 4). This is consistent with reduced protein abundance across all parts of the photosynthetic machinery including PSII, PSI, the Calvin cycle and the bicarbonate transport through Solute Carrier family 4 transporter, SLC4 (Nakajima *et al.*, 2013), as well as two carbonic anhydrases involved in the CO_2_ concentrating mechanism (Table I, Table S1). PSI was less widely affected than PSII, with a particularly strong reduction in levels of the PSI acceptor flavodoxin, while in the Calvin cycle there was a large reduction in levels of the key Calvin cycle enzyme, fructose bisphosphate aldolase FBAC5 (Gontero *et al.*, 2001; Erales *et al.*, 2008; Mininno *et al.*, 2012). In *P. tricornutum*, FBAC5 is located in the pyrenoid together with two fructose bisphosphate aldolases (Allen *et al.*, 2011). Finally, the decreased abundance of a bicarbonate transporter that is likely to supply the Calvin cycle with CO_2_ is reminiscent of the response of cyanobacteria to ppGpp accumulation (Hood *et al.*, 2016). Altogether these results indicate that altered PSII architecture and reduced photosynthetic capacity are highly conserved responses to ppGpp accumulation.

We also found evidence that ppGpp accumulation may slow cellular ageing in *P. tricornutum* by restricting growth, preventing chloroplast senescence, and blocking reserve compound accumulation. ppGpp accumulation caused a striking and sustained inhibition of growth (Fig. 2a) that is considerably stronger than that observed in plants (Maekawa *et al.*, 2015; Sugliani *et al.*, 2016). The arrest in growth was accompanied by the appearance of a single prominent LD, together with a slight increase in TAG (Fig. 5-6). The appearance of the prominent LD could be a consequence of a metabolic overshoot following growth arrest where the CO_2_ fixation rate exceeds requirements and the resulting carbon is diverted into the synthesis of reserve compounds. In the stationary phase, nutrients become limiting, and in wild type cells, this induces the synthesis of storage compounds and the degradation of photosynthetic pigments. However, we found that ppGpp over-accumulation prevented the degradation of chlorophyll and carotenoids (Fig. 3), and even led to increased levels of enzymes for chlorophyll biosynthesis (Table I). The storage compounds TAG and chrysolaminarin also did not increase in lines over-accumulating ppGpp, and we additionally observed higher levels of MGDG and SQDG, consistent with a more stable chloroplast (Fig. 5-6). Together these results suggest that ppGpp over-accumulation prevents cellular ageing and promotes a quiescent-like state. The anti-ageing effect of ppGpp in in *P. tricornutum* contrasts with the situation in plants where ppGpp accumulation leads to reduced chlorophyll levels, reduced chloroplast size and accelerated senescence (Maekawa *et al.*, 2015; Sugliani *et al.*, 2016). Together these results point to fundamental differences in (p)ppGpp signalling between multicellular and unicellular organisms (i.e. Arabidopsis versus *P. tricornutum*).

One of the most striking consequences of ppGpp accumulation in *P. tricornutum* was the strong induction of a wide range of chaperones and proteases (Table 1). Together these proteins can play an important role in protein protection by preventing protein aggregation and misfolding, and assisting with the refolding or destruction of denatured proteins (Saibil, 2013). Interestingly, the induced chaperones are likely distributed in the chloroplast, cytoplasm and mitochondria. The broad upregulation of protein quality control in response to ppGpp accumulation in *P. tricornutum* is reminiscent of the chloroplast unfolded protein response (cpUPR) in green algae and plants (Ramundo *et al.*, 2014; Ramundo & Rochaix, 2014; Llamas *et al.*, 2017; Perlaza *et al.*, 2019). The cpUPR is induced by the presence of misfolded proteins in the chloroplast, and leads to retrograde signalling and the accumulation of small heat-shock proteins, chaperones and proteases. The accumulation of ppGpp may therefore act as a signal for the presence of unfolded proteins, or alternatively could itself cause the accumulation of unfolded proteins for example via the inhibition of chloroplast gene expression as in plants (Yamburenko *et al.*, 2015; Sugliani *et al.*, 2016). Indeed, levels of deoxyxylulose-5-phosphate synthase, an enzyme of the chloroplastic isoprenoid pathway which accumulates in response to ppGpp in *P. tricornutum* (Table 1), are known to increase in response to defective chloroplast gene expression in Arabidopsis(Sauret-Gueto *et al.*, 2006; Llamas *et al.*, 2017). Interestingly, there are also parallels to the situation in bacteria. In *E. coli* for example, accumulation of (p)ppGpp during the stringent response leads to the induction of several chaperones (Grossman *et al.*, 1985).

ppGpp accumulation affects diatom physiology at multiple levels, and results in a quiescent-like state similar to the stringent response in bacteria that helps cells wait-out stress conditions (Fig. 8). Reduction of photosynthesis and growth arrest is a common response to most stresses in diatoms suggesting that ppGpp may play an important role in stress acclimation (Litchman *et al.*, 2003; Allen *et al.*, 2011; Brembu *et al.*, 2017). Other cell responses to ppGpp, such as the dramatic increase in HSP20, are similar to those found in response to dark stress (Bai *et al.*, 2016). However, despite these similarities, the effect of ppGpp accumulation is clearly different from those reported for nutrient stress. For example, under nitrogen starvation, there is a decrease in chlorophyll content and an increase in storage molecules like TAG (Longworth *et al.*, 2016). These comparisons suggest that ppGpp signalling may be an important component of a general diatom stress response pathway that acts in concertation with other acclimation pathways. Altogether, our findings highlight the importance of ppGpp as a fundamental regulator of chloroplast function across different domains of life, and lead to new questions about the molecular mechanisms and roles of (p)ppGpp signalling in photosynthetic eukaryotes.

## Acknowledgements

This work was supported by the Agence Nationale de la Recherche (ANR-15-CE05-0021-03, SignauxBioNRJ) and the IJPB Plant Observatory technological platform. The IJPB is supported by Saclay Plant Sciences-SPS (ANR-17-EUR-0007). We thank Julia Bartoli and Emmanuelle Bouveret for kindly providing 13C-ppGpp.

## Contributions

L.A., R.L., B.M., B.F. and B.G. conceived and planned the experiments. S. Citerne and G.M. performed the nucleotide quantification; S. Cuiné and Y. L-B. performed and analysed the lipidomics; L.A., C.P. and B.F. performed the remaining experiments. L.A., B.F. and B.G contributed to the interpretation of the results. L.A., B.F. and B.G. wrote the manuscript. All authors provided critical feedback and helped shape the research, analysis and manuscript.

## Supporting Information

**Figure S1. ppGpp levels in SYN and wild type cells under different conditions**.

**Figure S2. Growth curves of different SYN and SYN^D>G^ lines**.

**Figure S3. Pigment absorption spectra**.

**Figure S4. Photosynthetic parameters of SYN lines at different timepoints after induction**.

**Figure S5. Effect of ppGpp on polar lipid and fatty acid composition two days post induction**.

**Figure S6. Chrysolaminarin levels in SYN lines**.

**Figure S7. Protein profiles of SYN and controls two days after induction**.

**Figure S8. Volcano plot showing the changes in protein expression two days after SYN induction**.

**Table S1. List of primers used in this study**.

**Table S2. List of differentially expressed proteins in SYN lines versus controls**.

## References

Abdelkefi H, Sugliani M, Ke H, Harchouni S, Soubigou-Taconnat L, Citerne S, Mouille G, Fakhfakh H, Robaglia C, Field B. 2018. Guanosine tetraphosphate modulates salicylic acid signalling and the resistance of Arabidopsis thaliana to Turnip mosaic virus. Mol Plant Pathol 19(3): 634–646.

Abida H, Dolch L-J, Meï C, Villanova V, Conte M, Block MA, Finazzi G, Bastien O, Tirichine L, Bowler C, et al. 2015. Membrane Glycerolipid Remodeling Triggered by Nitrogen and Phosphorus Starvation in Phaeodactylum tricornutum. Plant Physiol 167(1): 118–136.

Allen AE, Dupont CL, Obornik M, Horak A, Nunes-Nesi A, McCrow JP, Zheng H, Johnson DA, Hu H, Fernie AR, et al. 2011. Evolution and metabolic significance of the urea cycle in photosynthetic diatoms. Nature 473(7346): 203–207.

Apt KE, Zaslavkaia L, Lippmeier JC, Lang M, Kilian O, Wetherbee R, Grossman AR, Kroth PG. 2002. In vivo characterization of diatom multipartite plastid targeting signals. J Cell Sci 115(Pt 21): 4061–4069.

Atkinson GC, Tenson T, Hauryliuk V. 2011. The RelA/SpoT homolog (RSH) superfamily: distribution and functional evolution of ppGpp synthetases and hydrolases across the tree of life. PLoS One 6(8): e23479.

Avilan L, Puppo C, Villain A, Bouveret E, Menand B, Field B, Gontero B. 2019. RSH enzyme diversity for (p)ppGpp metabolism in Phaeodactylum tricornutum and other diatoms. Sci Rep 9(1): 17682.

Bai X, Song H, Lavoie M, Zhu K, Su Y, Ye H, Chen S, Fu Z, Qian H. 2016. Proteomic analyses bring new insights into the effect of a dark stress on lipid biosynthesis in Phaeodactylum tricornutum. Sci Rep 6: 25494.

Bartoli J, Citerne S, Mouille G, Bouveret E, Field B. 2019. Quantification of guanosine tetraphosphate and other nucleotides in plants and algae using stable isotope-labelled internal standards. bioRxiv: 2019.2012.2013.875492.

Benoiston AS, Ibarbalz FM, Bittner L, Guidi L, Jahn O, Dutkiewicz S, Bowler C. 2017. The evolution of diatoms and their biogeochemical functions. Philos Trans R Soc Lond B Biol Sci 372(1728).

Bowler C, Allen AE, Badger JH, Grimwood J, Jabbari K, Kuo A, Maheswari U, Martens C, Maumus F, Otillar RP, et al. 2008. The Phaeodactylum genome reveals the evolutionary history of diatom genomes. Nature 456(7219): 239–244.

Brembu T, Muhlroth A, Alipanah L, Bones AM. 2017. The effects of phosphorus limitation on carbon metabolism in diatoms. Philos Trans R Soc Lond B Biol Sci 372(1728): 20160406.

Chu L, Ewe D, Rio Bartulos C, Kroth PG, Gruber A. 2016. Rapid induction of GFP expression by the nitrate reductase promoter in the diatom Phaeodactylum tricornutum. PeerJ 4: e2344.

Cox J, Hein MY, Luber CA, Paron I, Nagaraj N, Mann M. 2014. Accurate proteome-wide label-free quantification by delayed normalization and maximal peptide ratio extraction, termed MaxLFQ. Mol Cell Proteomics 13(9): 2513–2526.

Dorrell RG, Gile G, McCallum G, Meheust R, Bapteste EP, Klinger CM, Brillet-Gueguen L, Freeman KD, Richter DJ, Bowler C. 2017. Chimeric origins of ochrophytes and haptophytes revealed through an ancient plastid proteome. Elife 6.

Erales J, Avilan L, Lebreton S, Gontero B. 2008. Exploring CP12 binding proteins revealed aldolase as a new partner for the phosphoribulokinase/glyceraldehyde 3-phosphate dehydrogenase/CP12 complex--purification and kinetic characterization of this enzyme from Chlamydomonas reinhardtii. FEBS J 275(6): 1248–1259.

Fabris M, Matthijs M, Rombauts S, Vyverman W, Goossens A, Baart GJ. 2012. The metabolic blueprint of Phaeodactylum tricornutum reveals a eukaryotic Entner-Doudoroff glycolytic pathway. Plant J 70(6): 1004–1014.

Falciatore A, Casotti R, Leblanc C, Abrescia C, Bowler C. 1999. Transformation of Nonselectable Reporter Genes in Marine Diatoms. Mar Biotechnol (NY) 1(3): 239–251.

Falkowski P, Scholes RJ, Boyle E, Canadell J, Canfield D, Elser J, Gruber N, Hibbard K, Hogberg P, Linder S, et al. 2000. The global carbon cycle: a test of our knowledge of earth as a system. Science 290(5490): 291–296.

Field B. 2018. Green magic: regulation of the chloroplast stress response by (p)ppGpp in plants and algae. J Exp Bot 69(11): 2797–2807.

Flori S, Jouneau PH, Bailleul B, Gallet B, Estrozi LF, Moriscot C, Bastien O, Eicke S, Schober A, Bartulos CR, et al. 2017. Plastid thylakoid architecture optimizes photosynthesis in diatoms. Nat Commun 8: 15885.

Gontero B, Lebreton S, Graciet E 2001. Multienzyme complexes involved in the Benson-Calvin cycle and in fatty acid metabolism. In: McManus MT, Laing, W.A., Allan, A.C. ed. Annual Plant Reviews. Sheffield: Sheffield Academic Press, 120–150.

Granum E, Myklestad SM. 2002. A simple combined method for determination of β-1,3-glucan and cell wall polysaccharides in diatoms. Hydrobiologia 477(1): 155–161.

Greenspan P, Mayer EP, Fowler SD. 1985. Nile red: a selective fluorescent stain for intracellular lipid droplets. J Cell Biol 100(3): 965–973.

Grossman AD, Taylor WE, Burton ZF, Burgess RR, Gross CA. 1985. Stringent response in Escherichia coli induces expression of heat shock proteins. J Mol Biol 186(2): 357–365.

Gruber A, Kroth PG. 2017. Intracellular metabolic pathway distribution in diatoms and tools for genome-enabled experimental diatom research. Philos Trans R Soc Lond B Biol Sci 372(1728).

Gruber A, Rocap G, Kroth PG, Armbrust EV, Mock T. 2015. Plastid proteome prediction for diatoms and other algae with secondary plastids of the red lineage. Plant J 81(3): 519–528.

Guillard RR, Ryther JH. 1962. Studies of marine planktonic diatoms. I. Cyclotella nana Hustedt, and Detonula confervacea (cleve) Gran. Can J Microbiol 8(2): 229–239.

Harchouni S, Field B, Menand B. 2018. AC-202, a highly effective fluorophore for the visualization of lipid droplets in green algae and diatoms. Biotechnol Biofuels 11: 120.

Hauryliuk V, Atkinson GC, Murakami KS, Tenson T, Gerdes K. 2015. Recent functional insights into the role of (p)ppGpp in bacterial physiology. Nat Rev Microbiol 13(5): 298–309.

Honoki R, Ono S, Oikawa A, Saito K, Masuda S. 2018. Significance of accumulation of the alarmone (p)ppGpp in chloroplasts for controlling photosynthesis and metabolite balance during nitrogen starvation in Arabidopsis. Photosynth Res 135(1–3): 299–308.

Hood RD, Higgins SA, Flamholz A, Nichols RJ, Savage DF. 2016. The stringent response regulates adaptation to darkness in the cyanobacterium Synechococcus elongatus. Proc Natl Acad Sci U S A 113(33): E4867–4876.

Ihara Y, Ohta H, Masuda S. 2015. A highly sensitive quantification method for the accumulation of alarmone ppGpp in Arabidopsis thaliana using UPLC-ESI-qMS/MS. J Plant Res 128(3): 511–518.

Imamura S, Nomura Y, Takemura T, Pancha I, Taki K, Toguchi K, Tozawa Y, Tanaka K. 2018. The checkpoint kinase TOR (target of rapamycin) regulates expression of a nuclear-encoded chloroplast RelA-SpoT homolog (RSH) and modulates chloroplast ribosomal RNA synthesis in a unicellular red alga. Plant J 94(2): 327–339.

Ito D, Ihara Y, Nishihara H, Masuda S. 2017. Phylogenetic analysis of proteins involved in the stringent response in plant cells. J Plant Res 130(4): 625–634.

Jensen E, Clement R, Maberly SC, Gontero B. 2017. Regulation of the Calvin-Benson-Bassham cycle in the enigmatic diatoms: biochemical and evolutionary variations on an original theme. Philos Trans R Soc Lond B Biol Sci 372(1728).

Jeong JY, Yim HS, Ryu JY, Lee HS, Lee JH, Seen DS, Kang SG. 2012. One-step sequence- and ligation-independent cloning as a rapid and versatile cloning method for functional genomics studies. Appl Environ Microbiol 78(15): 5440–5443.

Joly N, Engl C, Jovanovic G, Huvet M, Toni T, Sheng X, Stumpf MP, Buck M. 2010. Managing membrane stress: the phage shock protein (Psp) response, from molecular mechanisms to physiology. FEMS Microbiol Rev 34(5): 797–827.

Krasny L, Gourse RL. 2004. An alternative strategy for bacterial ribosome synthesis: Bacillus subtilis rRNA transcription regulation. EMBO J 23(22): 4473–4483.

Kroth PG 2007. Genetic transformation:A Tool to Study Protein Targeting in Diatoms. In: Giezen vd ed. Protein Targeting Protocols. 2 ed. Methods in Molecular Biology. Totowa, NJ Humana Press Inc., 257–269.

Lavoie I, Hamilton PB, Morin S, Kim Tiam S, Kahlert M, Gonçalves S, Falasco E, Fortin C, Gontero B, Heudre D, et al. 2017. Diatom teratologies as biomarkers of contamination: Are all deformities ecologically meaningful? Ecological Indicators 82: 539–550.

Legeret B, Schulz-Raffelt M, Nguyen HM, Auroy P, Beisson F, Peltier G, Blanc G, Li-Beisson Y. 2016. Lipidomic and transcriptomic analyses of Chlamydomonas reinhardtii under heat stress unveil a direct route for the conversion of membrane lipids into storage lipids. Plant Cell Environ 39(4): 834–847.

Lepetit B, Goss R, Jakob T, Wilhelm C. 2012. Molecular dynamics of the diatom thylakoid membrane under different light conditions. Photosynth Res 111(1–2): 245–257.

Levitan O, Dinamarca J, Hochman G, Falkowski PG. 2014. Diatoms: a fossil fuel of the future. Trends Biotechnol 32(3): 117–124.

Litchman E, Steiner D, Bossard P. 2003. Photosynthetic and growth responses of three freshwater algae to phosphorus limitation and daylength. Freshwater Biology 48(12): 2141–2148.

Llamas E, Pulido P, Rodriguez-Concepcion M. 2017. Interference with plastome gene expression and Clp protease activity in Arabidopsis triggers a chloroplast unfolded protein response to restore protein homeostasis. PLoS Genet 13(9): e1007022.

Longworth J, Wu D, Huete-Ortega M, Wright PC, Vaidyanathan S. 2016. Proteome response of Phaeodactylum tricornutum, during lipid accumulation induced by nitrogen depletion. Algal Research 18: 213–224.

Maekawa M, Honoki R, Ihara Y, Sato R, Oikawa A, Kanno Y, Ohta H, Seo M, Saito K, Masuda S. 2015. Impact of the plastidial stringent response in plant growth and stress responses. Nat Plants 1: 15167.

Medlin LK. 2016. Evolution of the diatoms: major steps in their evolution and a review of the supporting molecular and morphological evidence. Phycologia 55(1): 79–103.

Mininno M, Brugiere S, Pautre V, Gilgen A, Ma S, Ferro M, Tardif M, Alban C, Ravanel S. 2012. Characterization of chloroplastic fructose 1,6-bisphosphate aldolases as lysine-methylated proteins in plants. J Biol Chem 287(25): 21034–21044.

Nagao R, Kato K, Suzuki T, Ifuku K, Uchiyama I, Kashino Y, Dohmae N, Akimoto S, Shen JR, Miyazaki N, et al. 2019. Structural basis for energy harvesting and dissipation in a diatom PSII-FCPII supercomplex. Nat Plants 5(8): 890–901.

Nakajima K, Tanaka A, Matsuda Y. 2013. SLC4 family transporters in a marine diatom directly pump bicarbonate from seawater. Proc Natl Acad Sci U S A 110(5): 1767–1772.

Nomura Y, Izumi A, Fukunaga Y, Kusumi K, Iba K, Watanabe S, Nakahira Y, Weber AP, Nozawa A, Tozawa Y. 2014. Diversity in Guanosine 3’,5’-Bisdiphosphate (ppGpp) Sensitivity Among Guanylate Kinases of Bacteria and Plants. J Biol Chem.

Ono S, Suzuki S, Ito D, Tagawa S, Shiina T, Masuda S. 2019. Plastidial (p)ppGpp synthesis by the Ca^2+^-dependent RelA-SpoT homolog regulates the adaptation of chloroplast gene expression to darkness in Arabidopsis. bioRxiv: 767004.

Perlaza K, Toutkoushian H, Boone M, Lam M, Iwai M, Jonikas MC, Walter P, Ramundo S. 2019. The Mars1 kinase confers photoprotection through signaling in the chloroplast unfolded protein response. eLife 8: e49577.

Pi X, Zhao S, Wang W, Liu D, Xu C, Han G, Kuang T, Sui SF, Shen JR. 2019. The pigment-protein network of a diatom photosystem II-light-harvesting antenna supercomplex. Science 365(6452).

Prioretti L, Avilan L, Carriere F, Montané M, Field B, Gregori G, Menand B, Gontero B. 2017. The inhibition of TOR in the model diatom Phaeodactylum tricornutum promotes a get-fat growth regime. Algal Research 26: 265–274.

Prioretti L, Carriere F, Field B, Avilan L, Montane MH, Menand B, Gontero B. 2019. Targeting TOR signaling for enhanced lipid productivity in algae. Biochimie.

Ramundo S, Casero D, Muhlhaus T, Hemme D, Sommer F, Crevecoeur M, Rahire M, Schroda M, Rusch J, Goodenough U, et al. 2014. Conditional Depletion of the Chlamydomonas Chloroplast ClpP Protease Activates Nuclear Genes Involved in Autophagy and Plastid Protein Quality Control. Plant Cell 26(5): 2201–2222.

Ramundo S, Rochaix JD. 2014. Chloroplast unfolded protein response, a new plastid stress signaling pathway? Plant Signal Behav 9(10): e972874.

Riesenberg D. 1985. A radioimmunoassay for (p)ppGpp and its application to Streptomyces hygroscopicus. J Basic Microbiol 25(2): 127–140.

Ritchie RJ. 2006. Consistent sets of spectrophotometric chlorophyll equations for acetone, methanol and ethanol solvents. Photosynth Res 89(1): 27–41.

Roding A, Boekema E, Buchel C. 2018. The structure of FCPb, a light-harvesting complex in the diatom Cyclotella meneghiniana. Photosynth Res 135(1–3): 203–211.

Rodionov DG, Ishiguro EE. 1995. Direct correlation between overproduction of guanosine 3’,5’-bispyrophosphate (ppGpp) and penicillin tolerance in Escherichia coli. J Bacteriol 177(15): 4224–4229.

Saibil H. 2013. Chaperone machines for protein folding, unfolding and disaggregation. Nat Rev Mol Cell Biol 14(10): 630–642.

Sambrook J, Fritsch E, Maniatis T. 1989. Molecular Cloning: a Laboratory Manual. : Cold Spring Harbor Laboratory Press, ColdSpring Harbor, NY.

Santin YG, Doan T, Lebrun R, Espinosa L, Journet L, Cascales E. 2018. In vivo TssA proximity labelling during type VI secretion biogenesis reveals TagA as a protein that stops and holds the sheath. Nat Microbiol 3(11): 1304–1313.

Sauret-Gueto S, Botella-Pavia P, Flores-Perez U, Martinez-Garcia JF, San Roman C, Leon P, Boronat A, Rodriguez-Concepcion M. 2006. Plastid cues posttranscriptionally regulate the accumulation of key enzymes of the methylerythritol phosphate pathway in Arabidopsis. Plant Physiol 141(1): 75–84.

Siaut M, Cuine S, Cagnon C, Fessler B, Nguyen M, Carrier P, Beyly A, Beisson F, Triantaphylides C, Li-Beisson Y, et al. 2011. Oil accumulation in the model green alga Chlamydomonas reinhardtii: characterization, variability between common laboratory strains and relationship with starch reserves. BMC Biotechnol 11: 7.

Siaut M, Heijde M, Mangogna M, Montsant A, Coesel S, Allen A, Manfredonia A, Falciatore A, Bowler C. 2007. Molecular toolbox for studying diatom biology in Phaeodactylum tricornutum. Gene 406(1–2): 23–35.

Steinchen W, Bange G. 2016. The magic dance of the alarmones (p)ppGpp. Mol Microbiol 101(4): 531–544.

Sugliani M, Abdelkefi H, Ke H, Bouveret E, Robaglia C, Caffarri S, Field B. 2016. An Ancient Bacterial Signaling Pathway Regulates Chloroplast Function to Influence Growth and Development in Arabidopsis. Plant Cell 28(3): 661–679.

Suzuki E, Suzuki R. 2013. Variation of Storage Polysaccharides in Phototrophic Microorganisms. Journal of Applied Glycoscience 60(1): 21–27.

Takahashi K, Kasai K, Ochi K. 2004. Identification of the bacterial alarmone guanosine 5’-diphosphate 3’-diphosphate (ppGpp) in plants. Proc Natl Acad Sci U S A 101(12): 4320–4324.

Tyanova S, Temu T, Sinitcyn P, Carlson A, Hein MY, Geiger T, Mann M, Cox J. 2016. The Perseus computational platform for comprehensive analysis of (prote)omics data. Nature Methods 13: 731.

Uthappa UT, Brahmkhatri V, Sriram G, Jung HY, Yu J, Kurkuri N, Aminabhavi TM, Altalhi T, Neelgund GM, Kurkuri MD. 2018. Nature engineered diatom biosilica as drug delivery systems. J Control Release 281: 70–83.

Vardi A, Bidle KD, Kwityn C, Hirsh DJ, Thompson SM, Callow JA, Falkowski P, Bowler C. 2008. A diatom gene regulating nitric-oxide signaling and susceptibility to diatom-derived aldehydes. Curr Biol 18(12): 895–899.

Wang LJ, Fan Y, Parsons RL, Hu GR, Zhang PY, Li FL. 2018. A Rapid Method for the Determination of Fucoxanthin in Diatom. Mar Drugs 16(1).

Yamburenko MV, Zubo YO, Borner T. 2015. Abscisic acid affects transcription of chloroplast genes via protein phosphatase 2C-dependent activation of nuclear genes: repression by guanosine-3’-5’-bisdiphosphate and activation by sigma factor 5. Plant J 82(6): 1030–1041.

