## Supporting Information for "ppGpp influences protein protection, growth and photosynthesis in *Phaeodactylum tricornutum*"

The following Supporting Information is available for this article:

**Fig. S1 ppGpp levels in SYN and wild type cells under different conditions.**

**Fig. S2 Growth curves of different SYN and SYN$^{D>G}$ lines**.

**Fig. S3 Pigment absorption spectra**.

**Fig. S4 Photosynthetic parameters of SYN lines at different timepoints after induction.**

**Fig. S5 Effect of ppGpp on polar lipid and fatty acid composition two days post induction**.

**Fig. S6 Chrysolaminarin levels in SYN lines**.

**Fig. S7 Protein profiles of SYN and controls two days after induction**.

**Fig. S8 Volcano plot showing the changes in protein expression two days after SYN induction**.

**Table S1 List of different primers used in this study.**

**Table S2 List of differentially expressed proteins in SYN lines versus controls.**

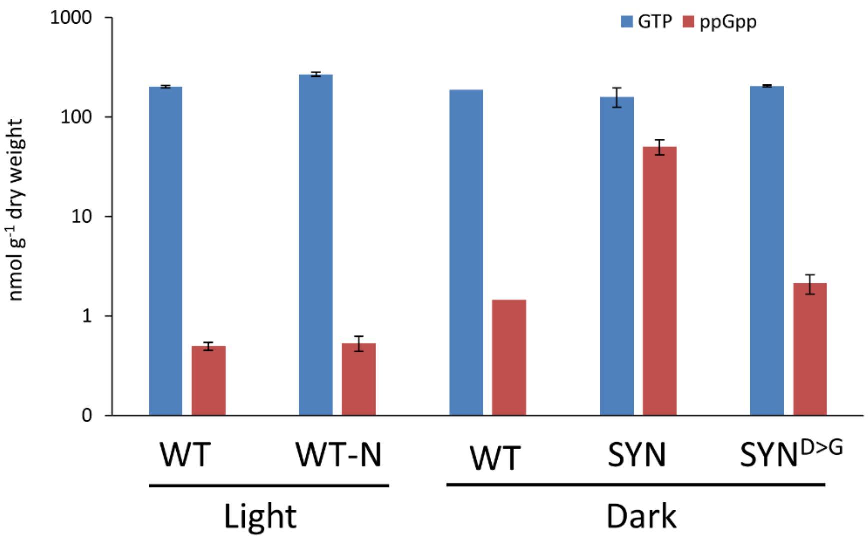


**Fig. S1 . ppGpp levels in SYN and wild type cells under different conditions.** ppGpp and GTP levels in SYN (±SE, n=5 independent lines), $SYN^{D>G}$ (±SE, n =4 independent lines) and WT (1 replicate) 21 days post induction with incubation in the dark, and in WT cultured in the light in standard f/2-$NO_3$ medium or f/2 medium without nitrogen (-N) (±SE, n= 3 biological replicates).

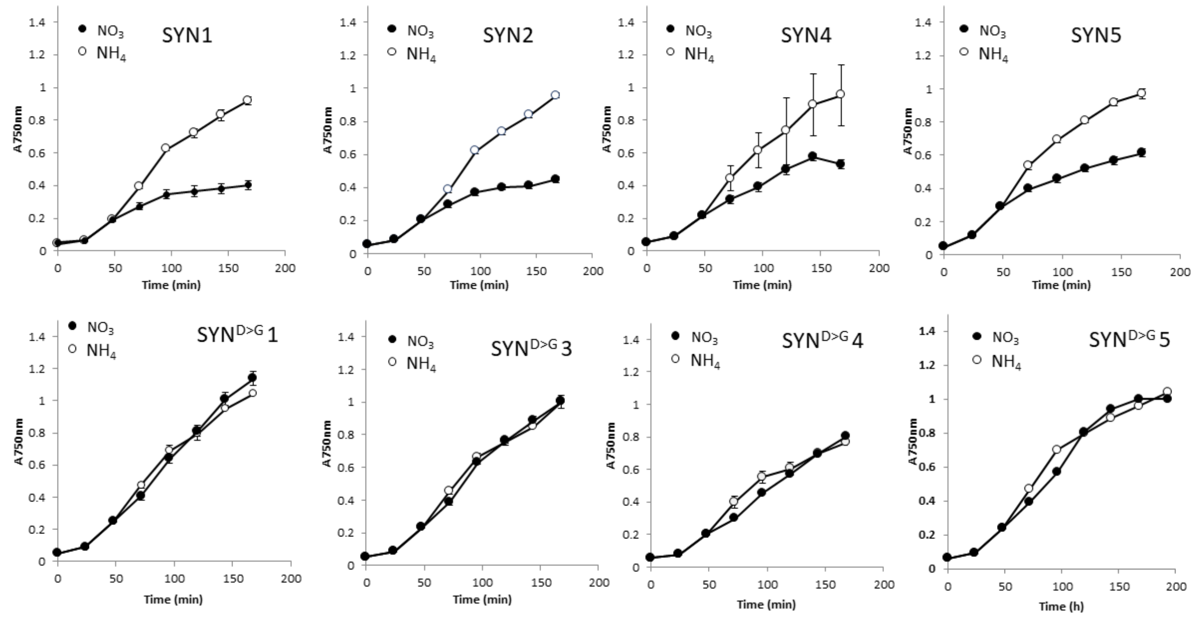


**Fig. S2 Growth curves of different SYN and SYN$^{D>G}$ lines**. Cells grown in f/2-$NH_4$ were transferred after two days to either f/2-$NO_3$ for induction or f/2-$NH_4$ for the non-induced control.

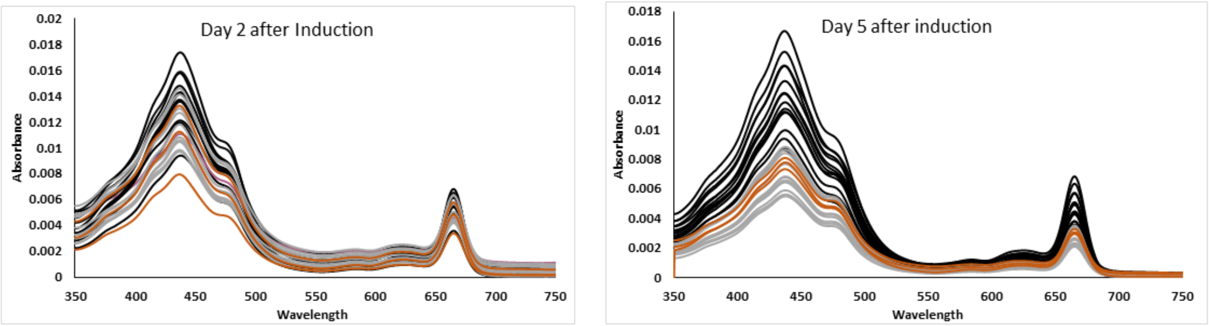


**Fig. S3 Pigment absorption spectra**. Pigments from SYN (black), $SYN^{D>G}$ (gray) and WT (brown) cells were extracted with ethanol. Cells were grown in f/2-$NH_4$ and transferred to f/2-$NO_3$ for induction and harvested two and five days post induction.

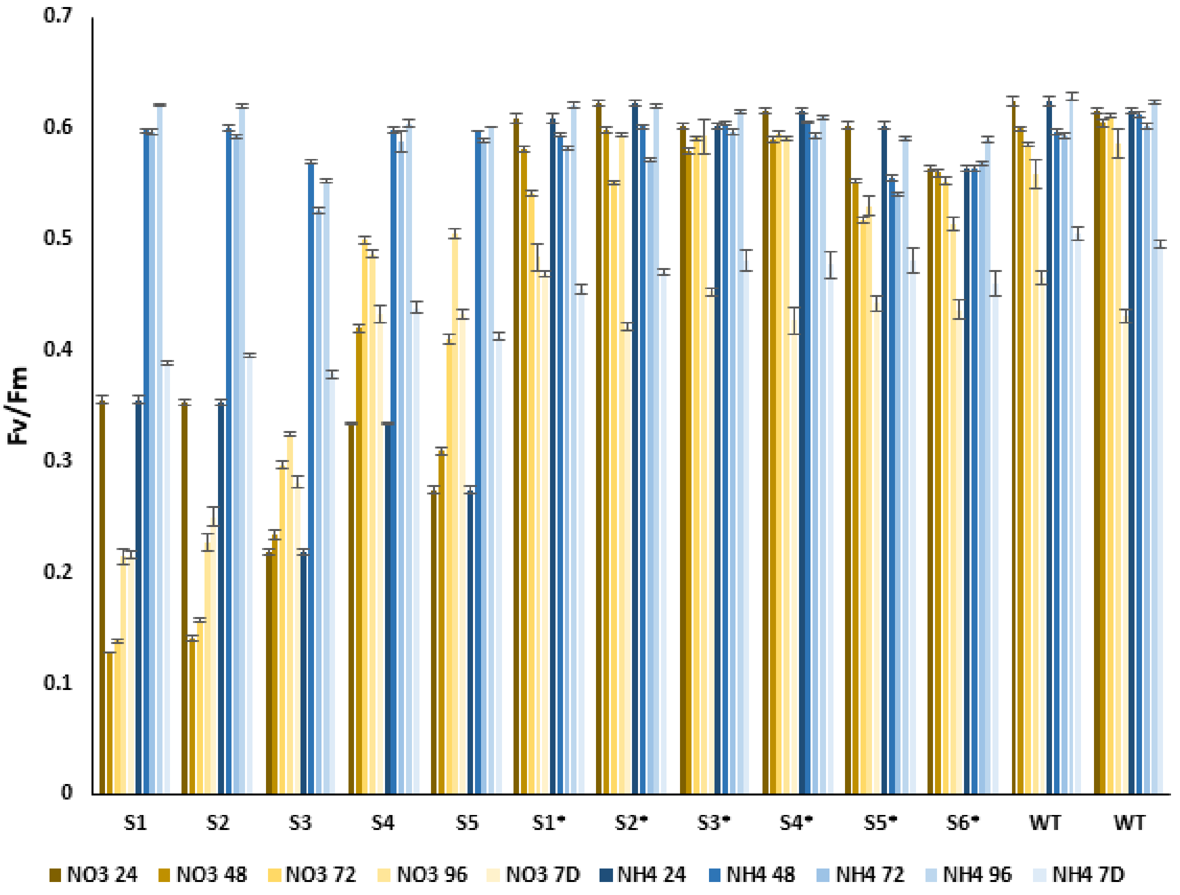


**Fig. S4 Photosynthetic parameters of SYN lines at different timepoints after induction.** The maximal efficiency of the photosystem II ($F_v/F_m$) was measured at 24, 48, 72, 96 hours and 7 days post induction with f/2-$NO_3$, in five SYN lines, six $SYN^{D>G}$ lines and WT. Controls were cultures grown in in f/2-$NH_4$. Averages are shown, ± SE, n=7.

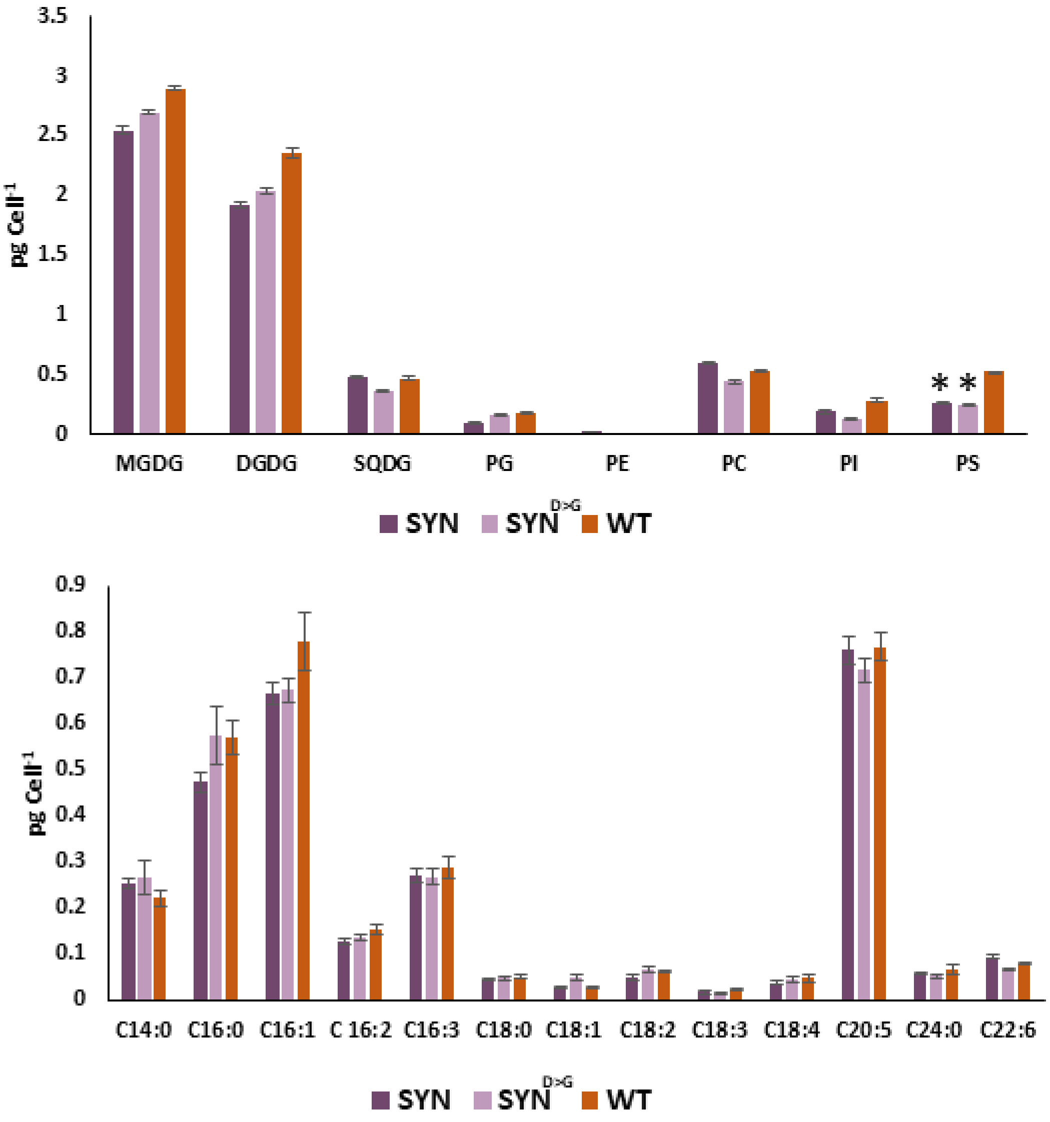


**Fig. S5 Effect of ppGpp on polar lipid and fatty acid composition two days post induction**. **(a)** Polar lipids and **(b)** fatty acid levels were determined two days post induction. MGDG, monogalactosyldiacylglycerol; DGDG, digalactosyldiacylglycerol; SQDG, sulfoquinovosyldiacylglycerol; PG, phosphatidylglycerol; PE, phosphatidylethanolamine; PC, phosphatidylcholine; PI, phosphatidylinositol and PS, phosphatidylserine. Data are means ± SE of five biological replicates, analysed by ANOVA using Dunnett post-hoc test versus WT control, *$P<0.05$, **$P < 0.01$.

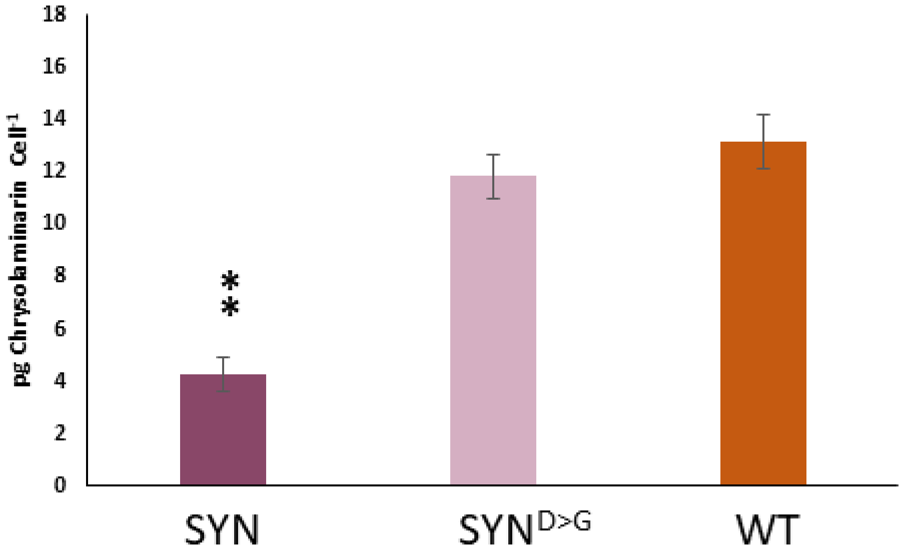


**Fig. S6 Chrysolaminarin levels in SYN lines**. Chrysolaminarin levels were determined five days post induction in cell pellets containing 2 x $10^7$ cells. Data are means ± SE of five biological replicates, analysed by ANOVA using Dunnett post-hoc test versus WT control, $**P < 0.01$.

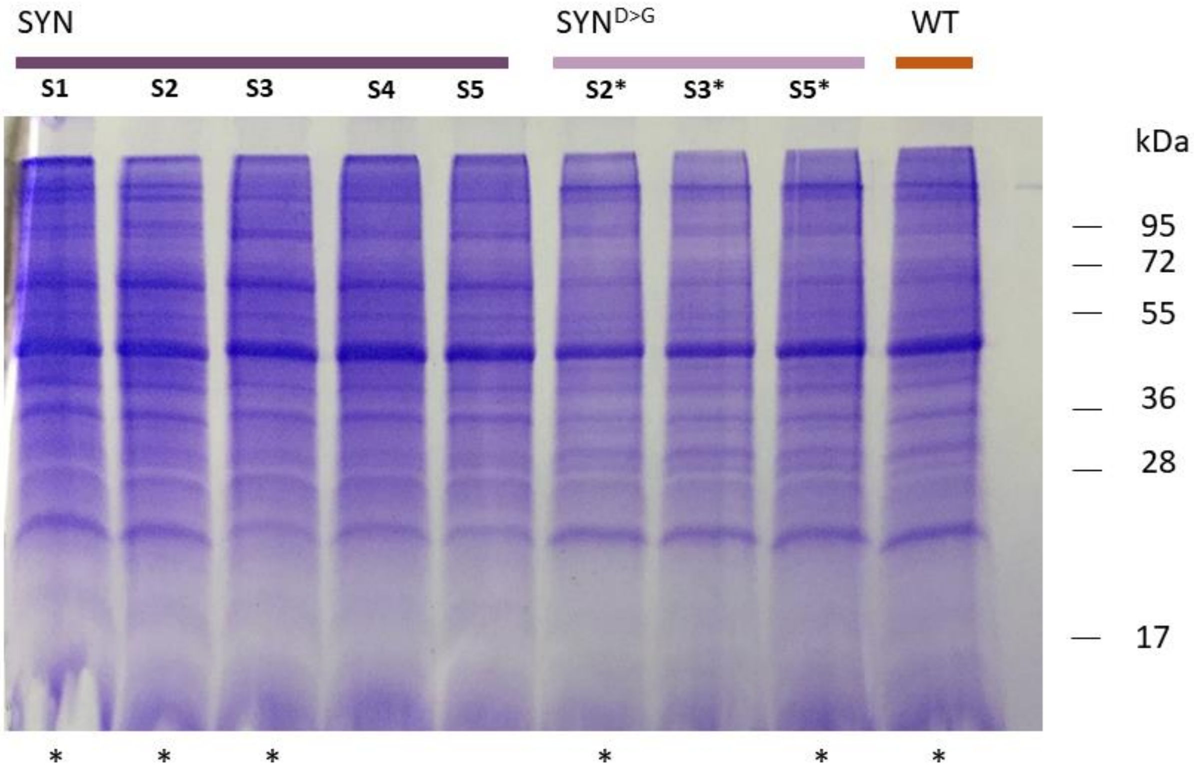


**Fig. S7 Protein profiles of SYN and controls two days after induction**. Protein extracts (50 µg protein) from several induced SYN and SYN$^{D>G}$ lines and the WT, were separated by SDS-PAGE and stained with Coomassie Brilliant Blu(e) Stars below the gel indicate the samples used for proteomic analysis.

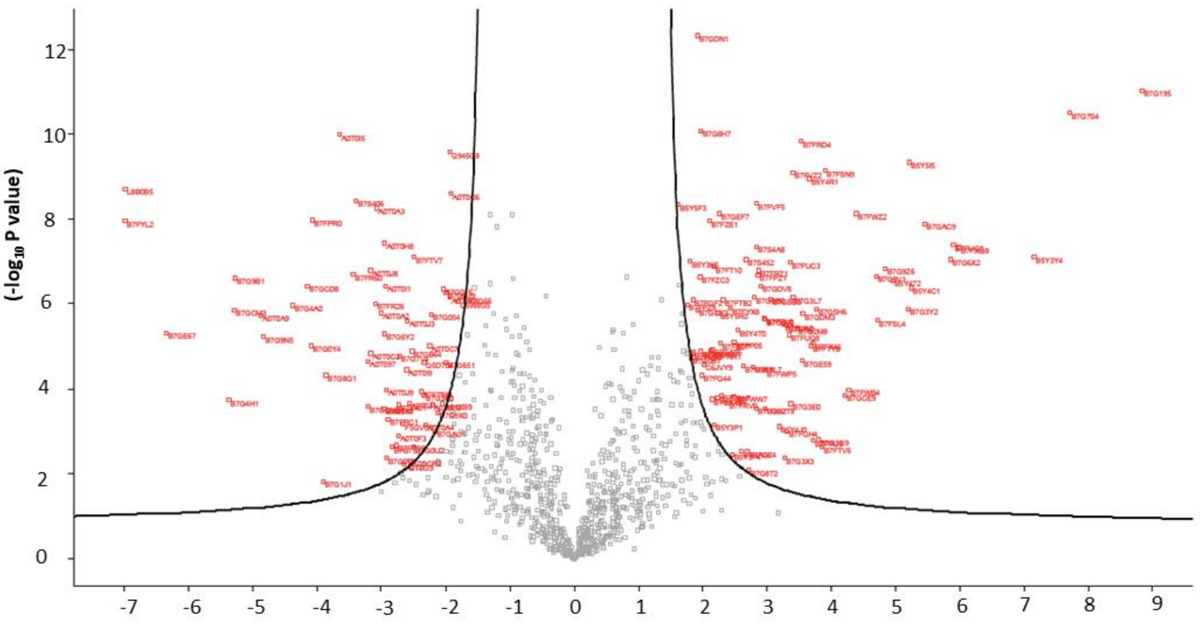


Fig. S8 -$\log_2$ Fold change (ppGpp-control)

**Fig. S8 Volcano plot showing the changes in protein expression two days after SYN induction**. The plot highlights the differentially accumulating proteins in SYN lines. The x-axis shows $\log_2$ fold change of control ($SYN^{D>G}$ and WT) versus SYN. The y-axis shows the log10 p value. Proteins with a p value < 0.1 and a fold change of 4 are highlighted in red.

**Table S1. List of primers used in this study**

| Name | Sequence (5'-3') | Experiment |
|---|---|---|
| ATPc-up | CACTTGTGCGAACGGAATTCAAGATGAGAT CCTTTTGCATCGC | Amplification of chloroplastic gamma ATP synthase bipartite targeting peptide and cloning |
| ATPc-Syn-low | TTACCGCAAC CATGACAATCGTTGCTTTACG | Amplification of chloroplastic gamma ATP synthase bipartite targeting peptide and cloning |
| Syn-up | GATTGTCATG GTTGCGGTAAGAAGTGCACA | Amplification of SYN and cloning |
| Syn- low | CTTAAAGTAAATTGAAGCTTTTAATGGTGAT GGTGATGGT | Amplification of SYN and cloning |
| ATP-Syn-up | ATGAGATCCTTTTGCATCGCAGC | Screen PCR |
| ATP-Syn-low | AATGGTGATGGTGATGGTGTCCA | Screen PCR |
| Sh-ble-up | TCGAGTTCTGGACCGACCGGCT | Screen PCR |
| Sh-ble-low | ACGAAGTGCACGCAGTTGCCGG | Screen PCR |
